# Seizures, behavioral deficits and adverse drug responses in two new genetic mouse models of *HCN1* epileptic encephalopathy

**DOI:** 10.1101/2021.08.16.456452

**Authors:** Andrea Merseburg, Jacquelin Kasemir, Eric W. Buss, Felix Leroy, Tobias Bock, Alessandro Porro, Anastasia Barnett, Simon E. Tröder, Birgit Engeland, Malte Stockebrand, Anna Moroni, Steven A Siegelbaum, Dirk Isbrandt, Bina Santoro

## Abstract

*De novo* mutations in voltage- and ligand-gated channels have been associated with an increasing number of cases of developmental and epileptic encephalopathies, which often fail to respond to classic antiseizure medications. Here, we examine two knock-in mouse models replicating *de novo* mutations in the *HCN1* voltage-gated channel gene, p.G391D and p.M153I (*Hcn1^G380D/+^* and *Hcn1^M142I/+^* in mouse), associated with severe drug-resistant neonatal- and childhood-onset epilepsy, respectively. Heterozygous mice from both lines displayed spontaneous generalized tonic-clonic seizures. *Hcn1*^G380D/+^ animals had an overall more severe phenotype, with pronounced alterations in the levels and distribution of HCN1 protein, including disrupted targeting to the axon terminals of basket cell interneurons. In line with clinical reports from *HCN1* patients, administration of the antiepileptic Na^+^ channel antagonists lamotrigine and phenytoin resulted in the paradoxical induction of seizures in both lines, consistent with an effect to further impair inhibitory neuron function. We also show that these variants can render HCN1 channels unresponsive to classic antagonists, indicating the need to screen mutated channels to identify novel compounds with diverse mechanism of action. Our results underscore the need to tailor effective therapies for specific channel gene variants, and how strongly validated animal models may provide an invaluable tool towards reaching this objective.

## Introduction

Developmental and epileptic encephalopathies (DEE) are a devastating group of diseases, often with poor response to pharmacological treatment, resulting in a lifelong burden of seizures, developmental delay, and intellectual disability. Since the original discovery over twenty years ago that *de novo* mutations in voltage-gated ion channels can directly cause early childhood epilepsy syndromes (Singh et al., 1998), the number of genes and gene variants associated with DEE has grown exponentially. Current estimates indicate that ∼30% of DEE patients carry at least one pathogenic variant in the top 100 known gene candidates for the disease, with about a third of the affected genes falling into the category of voltage- and ligand-gated ion channels (Noebels, 2017; Oyrer et al., 2018; Wang and Frankel, 2021). Despite the wealth of genetic data available and recent efforts to model the effects of the mutations in genetically modified mice, we understand very little about the mechanisms that underlie the brain and neuronal circuit alterations responsible for seizures, or the reasons for their drug resistance.

Here we focus on seizures and the pharmacological response profile associated with mutations in the *HCN1* gene, which encodes a hyperpolarization-activated cyclic nucleotide-regulated non-selective cation channel expressed prominently in brain (Santoro and Shah, 2020). To date a total of 40 different missense variants in the *HCN1* gene have been reported in patients affected by epilepsy and/or neurodevelopmental disorders (source: HGMD Professional 2020.4). Among these, at least 12 different *de novo HCN1* variants have been linked to DEE (Nava et al., 2014; Butler et al., 2017; Marini et al., 2018; Wang et al., 2019).

The functional role of HCN1 channels is tied closely to their distinct subcellular localization in the two classes of neurons where HCN1 protein is primarily expressed, pyramidal neurons in cortex and hippocampus, and parvalbumin positive (PV+) interneurons across the brain (Notomi and Shigemoto, 2004). Within these neuronal classes, the subcellular localization of HCN1 channel subunits is tightly regulated. In excitatory (pyramidal) neurons, the channel is targeted to the distal portion of the apical dendrites, where it constrains the dendritic integration of excitatory postsynaptic potentials, limiting excitability (Tsay et al., 2007; George et al., 2009). In inhibitory (PV+) neurons, HCN1 channels are localized exclusively to axonal terminals, where they facilitate rapid action potential propagation (Roth and Hu, 2020) and GABA release (Southan et al., 2000; Aponte et al., 2006).

In this study we examine two new genetic mouse models that reproduce *de novo HCN1* variants previously shown to be associated with severe forms of neonatal- and childhood-onset epilepsy (Parrini et al., 2017; Marini et al., 2018; Fernández-Marmiesse et al., 2019). A prior attempt at modeling *HCN1*-linked DEE in mice (Bleakley et al., 2021) resulted in a relatively mild phenotype, wherein spontaneous seizures could not be systematically documented, despite the presence of epileptiform spikes on electrocorticography recordings (ECoG). This prompts the question whether there may be some general limitation to modeling *HCN1*-linked DEE in mice. To examine this question and probe further whether and how *HCN1* mutations may give rise to DEE, we generated mouse models for two other *HCN1* variants linked to DEE in humans.

We examined variants c.1172G>A/p.Gly391Asp and c.459G>C/p.Met153Ile based on three criteria: severity of disease; occurrence in at least two independent patients with similar epilepsy phenotypes; and differing biophysical effects on channel function when tested in heterologous expression systems (Marini et al., 2018). Moreover, clinical reports suggest that seizures are poorly controlled with standard antiseizure medications, and that certain anticonvulsants in fact exacerbate seizures in these patients (Marini et al., 2018). Our results show that both mutations lead to spontaneous seizures in mice, reproduce the paradoxical pharmacological responses seen in patients, and provide new insights into the mechanistic basis for the limitations of current drug therapies.

## Results

Pathological mutations orthologous to human *HCN1* variants p.G391D and p.M153I were generated using the CRISPR/Cas9 approach in the *Hcn1* gene of inbred C57BL/6J mice (see Materials and Methods; Tröder et al., 2018). The resulting lines, *Hcn1^G380D/+^* and *Hcn1^M142I/+^*, were maintained using a male heterozygote x female wildtype (WT) breeding scheme and heterozygous mutant animals were compared to WT littermate controls.

### General phenotypic features and gross brain anatomy

No overt differences were noted in pup development, except for consistently smaller body weights at weaning in *Hcn1^G380D/+^* mice of both sexes (Figure 1A). Smaller body weights persisted for the life of *Hcn1^G380D/+^* animals, and were accompanied by an ∼12% reduction in adult brain weight with gross brain anatomy otherwise normal (Figure 1B). Brain area measurements showed a similar reduction. Thus, both cerebellum and brainstem areas were significantly smaller in *Hcn1^G380D/+^* mice compared to WT (Figure 1 – Figure supplement 1) with the cerebellum more affected than the brainstem, consistent with the higher expression of HCN1 protein in this region (Notomi and Shigemoto, 2004). Reduced mean body weight was also observed in *Hcn1^M142I/+^* males, but no decrease in brain size was noted in either sex for this line (Figure 1A,B).

**Figure 1:**
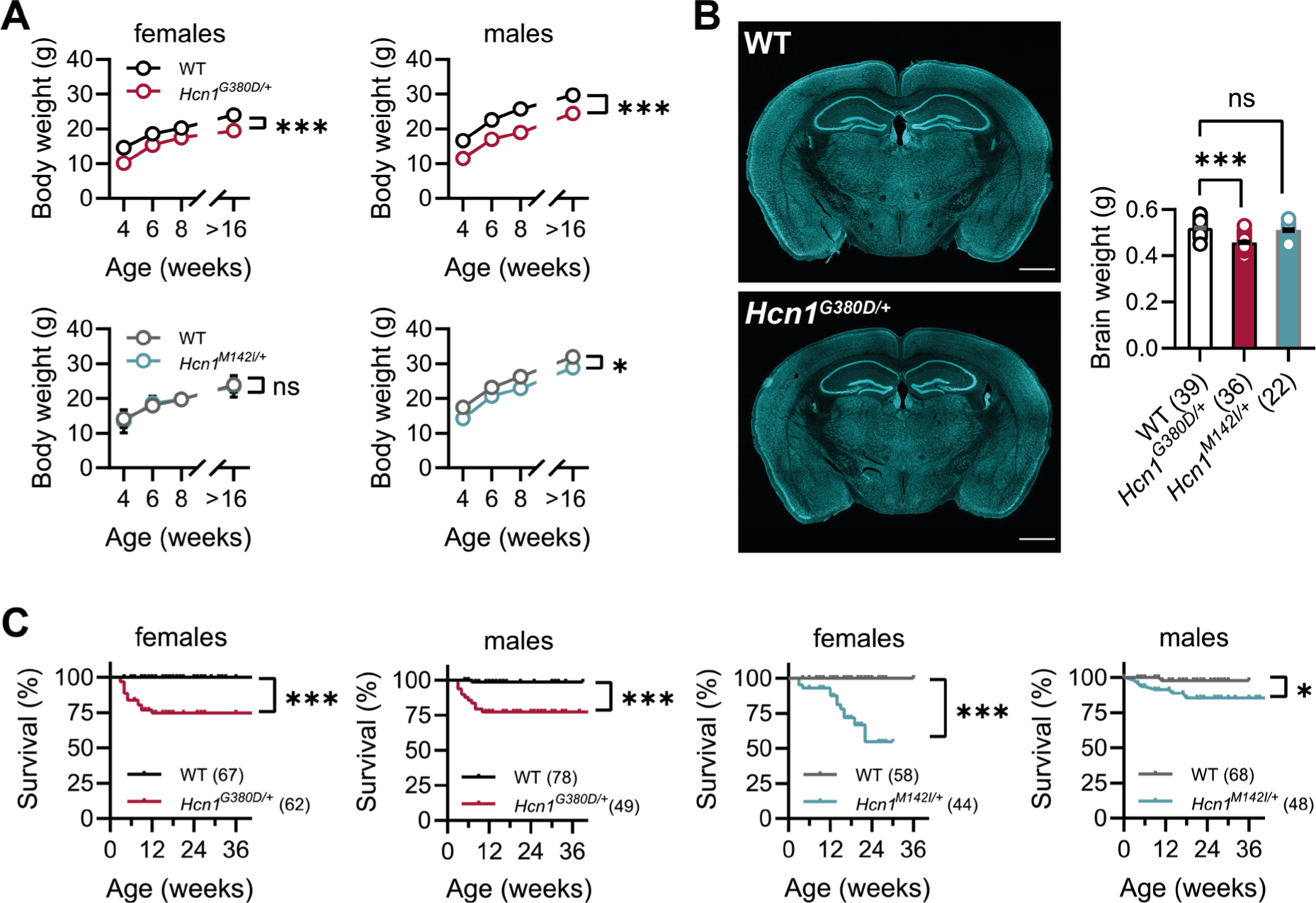
Body weight, survival and general brain morphology in *Hcn1^G380D/+^* and *Hcn1^M142I/+^* mice. A) Reduced body weight post-weaning and through adult life in *Hcn1^G380D/+^* mice of both sexes, as well as in *Hcn1^M142I/+^* males (ns = not significant, * *P* < 0.05, *** *P* < 0.001; effect of *genotype*, mixed-effects model having *age* x *genotype* as grouping factors, G380D: WT females n=11-29, *Hcn1^G380D/+^* females n=9-23, WT males n=10-38, *Hcn1^G380D/+^* males n=9-31; M142I: WT females n=11-28, *Hcn1^M142I/+^* females n=5-17, WT males n=5-16, *Hcn1^M142I/+^* males n=9- 20). B) Brain weight in WT, *Hcn1^G380D/+^* and *Hcn1^M142I/+^* mice. Fluorescent Nissl stain of mid- coronal brain section from WT and *Hcn1^G380D/+^* mice shown on the left (scale bar = 1200 μm). Brain weight shown on the right (WT 0.52 ± 0.01 g; *Hcn1^G380D/+^* 0.46 ± 0.01 g; *Hcn1^M142I/+^* 0.51 ± 0.01 g, ns = not significant; *** *P* < 0.001, after one-way ANOVA and *post hoc* Holm-Šídák’s multiple comparisons test). Brain weights were measured on PFA-fixed tissue, after removal of both olfactory bulbs. Data represent mean ± S.E.M. C) Kaplan-Meier survival curves show increased mortality in both *Hcn1^G380D/+^* and *Hcn1^M142I/+^* mice, with a marked sex difference in the case of *Hcn1^M142I/+^* animals. Each dot represents a death event in the colony, scored as 0 if resulting from experimental use and 1 if due to sudden or unprovoked death (survival at 36 weeks: *Hcn1^G380D/+^* females = 75%, *Hcn1^G380D/+^* males = 77%, *Hcn1^M142I/+^* females = 55%, *Hcn1^M142I/+^* males = 85%, * *P* < 0.05, *** *P* < 0.001 after log-rank Mantel-Cox test).

Survival curves revealed a significant number of premature deaths among mutant animals in both lines (Figure 1C). In many instances, animals were found dead in the cage without having shown prior signs of distress, suggesting sudden death. The survival curves revealed some notable differences in the pattern of death observed between the two lines, both with regards to timing and sex specificity. Thus, while most deaths among *Hcn1^G380D/+^* animals occurred before the age of three months, with no difference between the sexes (Figure 1C, left), there was a clear split in *Hcn1^M142I/+^* animals with females more affected than males and most deaths occurring at an age older than three months (Figure 1C, right). These findings suggest important differences in the effects of the two variants on brain function and physiology, perhaps reflective of the divergent effects of the mutations on channel biophysical properties (Marini et al., 2018) and HCN1 protein expression (see below).

Given the high expression of HCN1 subunits in cerebellum, both in Purkinje neurons and basket cell axon terminals (Southan et al., 2000; Luján et al., 2005; Rinaldi et al., 2013), as well as the motor phenotype observed in HCN1 global knockout animals (Nolan et al., 2003; Massella et al., 2009), we tested for basic locomotion in an open field setting (Figure 2A) and assessed gait parameters (Figure 2D). Mutant animals in both lines showed hyperactivity in the open field, with significant increases in running speed and distance moved, but a reduction in the center *versus* border zone occupation ratio, potentially indicative of increased anxiety (Figure 2B,C and Figure 2 – Figure supplement 1). Gait analysis reflected the overall higher running speed of *Hcn1^G380D/+^* and *Hcn1^M142I/+^* animals, compared to WT littermates (Figure 2E). When runs of similar speed were compared, significant changes in the base of support (BOS) were observed in *Hcn1^G380D/+^* animals, with the front paws’ BOS being wider at both speed ranges (Figure 2F), and the hind paws’ BOS narrower at higher running speeds (Figure 2 – Figure supplement 2). No BOS alterations were noted in *Hcn1^M142I/+^* animals, or any of the other parameters measured (Figure 2F and Figure 2 – figure supplements 2 and 3).

**Figure 2:**
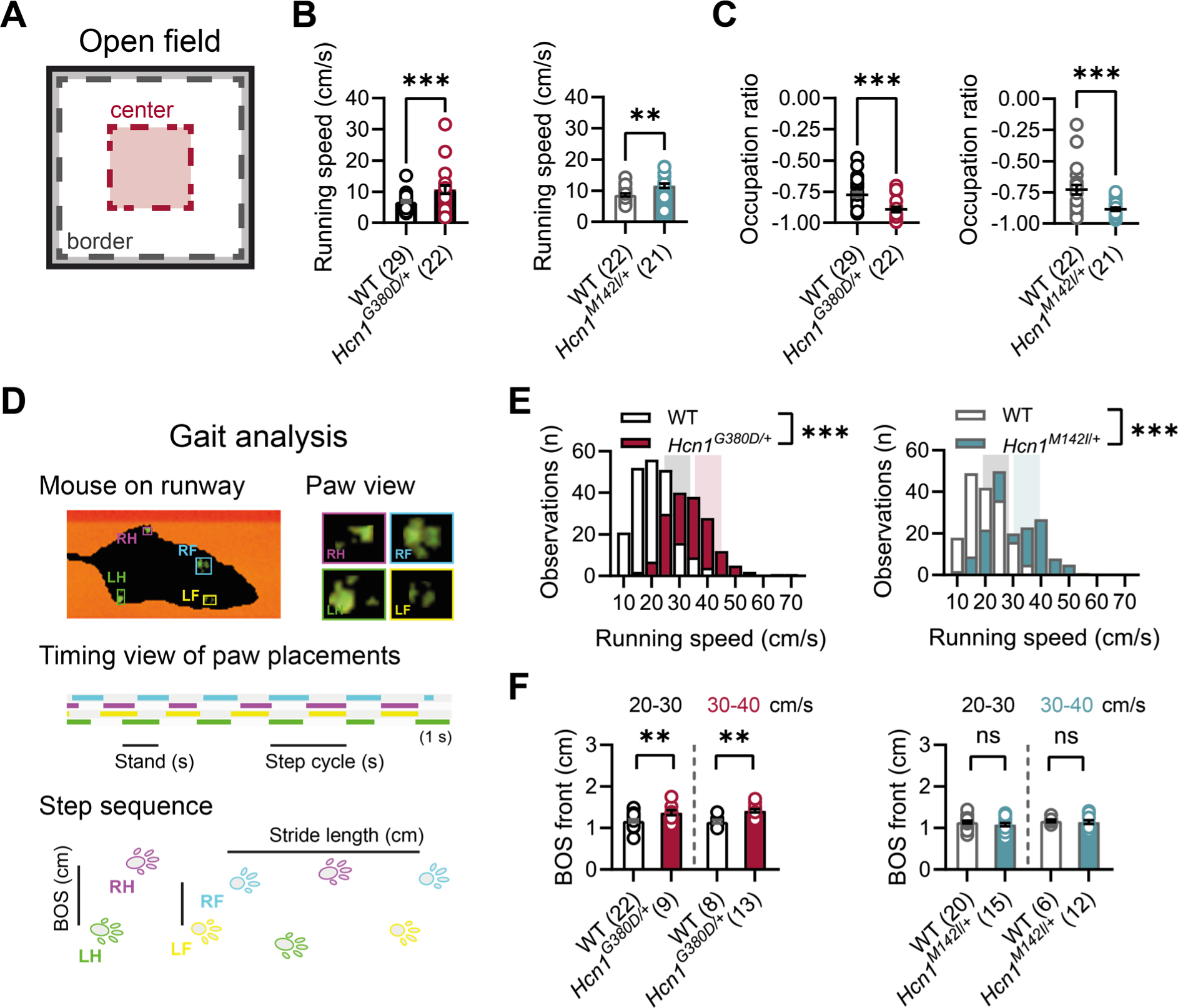
Baseline exploratory behavior and gait analysis in *Hcn1^G380D/+^* and *Hcn1^M142I/+^* mice. A) Schematic representation of the open field arena, with dashed lines depicting the imaginary border (grey) and center zones (red). Occupation ratio calculated as (Center- Border)/(Center+Border), with values < 0 representing a preference for the border over the center zone. Mean running speed was significantly increased (B) and mean occupation ratio decreased (C) in both *Hcn1^G380D/+^* and *Hcn1^M142I/+^* mice compared to WT littermates (** *P* < 0.01; *** *P* < 0.001, Mann Whitney *U* test.) C) Representation of the Catwalk gait analysis system, depicting the mouse on the runway (top left), the paw print view (top right), and below the timing view of the consecutive paw placements and step sequence, with the base of support (BOS) representing the distance between the hind and the front paws, respectively. E) Distribution of running speeds of *Hcn1^G380D/+^* or *Hcn1^M142I/+^* mice compared to WT littermates (*** *P* < 0.001, Mann Whitney *U* test). Multiple runs were performed for each animal (G380D: WT 30 mice, n=211 runs, mean number of runs per animal = 6.1 ± 0.4; *Hcn1^G380D/+^* 21 mice, n=166 runs, mean number of runs per animal = 7.7 ± 0.4. M142I: WT 22 mice, n=166 runs, mean number of runs per animal = 7.6 ± 0.2; *Hcn1^M142I/+^* 21 mice, n=167 runs, mean number of runs per animal = 8.0 ± 0.1). Shaded areas highlight speed range of 20-30 cm/sec (grey) and 30-40 cm/s (color). F) BOS of front paws was significantly increased in *Hcn1^G380D/+^* at both speed ranges, but not in *Hcn1^M142I/+^* mice (** *P* < 0.01, ns = not significant, Student’s *t* test). Data in F represent mean ± S.E.M. for indicated number of animals. Only runs in the selected speed range were counted and BOS values averaged per animal (see Figure 2 – source data 1 and 2 for numerical values).

In summary, these results reveal a complex effect of the two variants on phenotype, with an overall more severe impact on the general fitness of *Hcn1^G380D/+^* compared to *Hcn1^M142I/+^* animals, reflected in smaller body size, reduced brain size, as well as altered gait and locomotion.

### Hcn1^G380D/+^ and Hcn1^M142I/+^ mice show spontaneous convulsive seizures

The sudden death phenotype observed in the two lines, along with the epilepsy syndromes seen in G391D and M153I HCN1 patients, suggests the animals may be undergoing generalized tonic-clonic seizures (GTCS), which in mice often result in death when escalating into tonic hindlimb extension. One such death event was indeed observed in a female from the *Hcn1^M142I/+^* line, and several convulsive seizure events were witnessed in mice from both lines during routine cage inspections. We therefore set out to rigorously quantify the occurrence, frequency and severity of seizures in *Hcn1^G380D/+^* and *Hcn1^M142I/+^* mice by using chronic video-ECoG recordings for periods of 1-2 weeks at a time (Figure 3). Adult mice from both lines and sexes were found to display spontaneous GTCS’s with distinct electrographic signature on ECoG traces (Figure 3A,C) and behavioral manifestation on video recordings (Videos 1 and 2).

**Figure 3:**
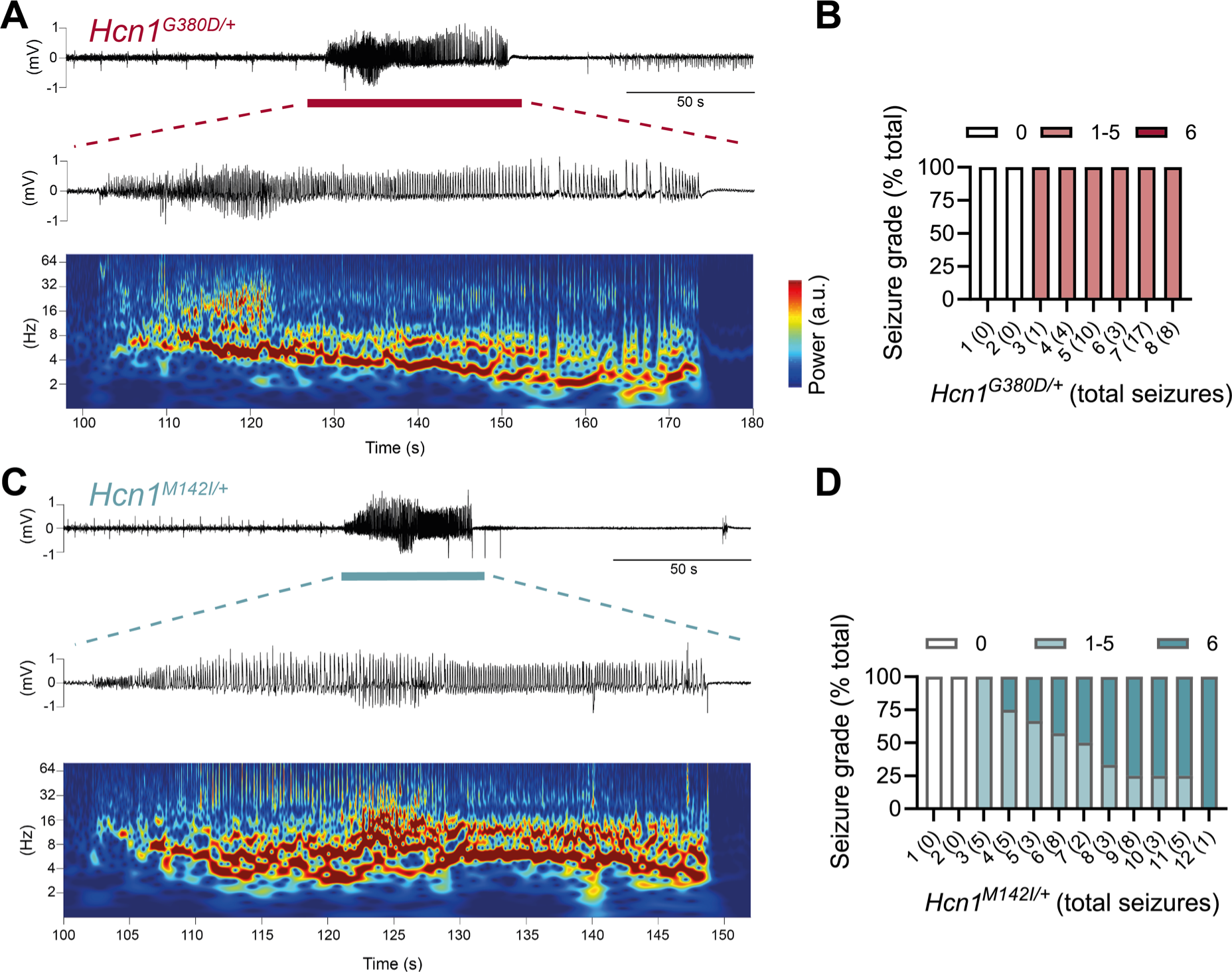
Spontaneous convulsive seizures in *Hcn1^G380D/+^* and *Hcn1^M142I/+^* mice. A) Example of a spontaneous seizure recorded in an *Hcn1^G380D/+^* mouse. Upper trace shows ECoG-signal of a behaviorally noted grade 4 seizure (red bar), with expanded trace and corresponding time-frequency spectrogram shown below. B) Highest seizure grade reached (plotted as percentage of total seizure number) for all individual *Hcn1^G380D/+^* mice examined, with total number of seizure events observed during the recording period shown in parentheses. 2/8 animals did not exhibit spontaneous seizures during the recording period of 12 ± 4 days. Of the 6 individuals with seizures, only 1 animal reached seizures with a grade of 6 on a modified Racine’s scale (see Materials and Methods). Mean seizure frequency for *Hcn1^G380D/+^* animals was 0.33 ± 0.08/day. C) Same as in (A) showing a grade 6 seizure in an *Hcn1^M142I/+^* mouse. D) Highest seizure grade reached as compared to total seizure event number shown for the 12 individual *Hcn1^M142I/+^* mice. 2/12 animals did not exhibit spontaneous seizures during the recording period. Of the 10 individuals with seizures, 9 animals reached a seizure grade of 6, indicating an overall higher severity of the spontaneous seizures in *Hcn1^M142I/+^* animals. Mean seizure frequency for *Hcn1^M142I/+^* animals was 0.42 ± 0.07/day (females, 0.40 ± 0.08/day; males, 0.45 ± 0.11/day; *P* = 0.7).

We recorded from a total of n = 8 *Hcn1^G380D/+^* animals (2M, 6F) 56-91 days old to avoid selecting against animals with early death; and a total of n = 12 *Hcn1^M142I/+^* animals (6M, 6F) 80-130 days old to capture the period during which sudden death is most frequently seen in females. Notably, despite the higher occurrence of death in females, seizures were observed with similar frequency in males and females from the *Hcn1^M142I/+^* line (Figure 3D). Seizure events were comparatively infrequent, ranging from 0.6 - 4.1 seizures per week in *Hcn1^G380D/+^* mice (with 2/8 individuals failing to show seizures during the time recorded) and from 0.6 - 5.1 seizures per week in *Hcn1^M142I/+^* mice (with 2/12 individuals displaying no seizures during the recorded period). Despite the overall more severe phenotype of *Hcn1^G380D/+^* animals, based on body weight, brain size and general appearance, when rated on a modified Racine scale (see Materials and Methods) we found that animals from the *Hcn1^G380D/+^* line had on average lower maximum grade seizures compared to *Hcn1^M142I/+^* animals (Figure 3B,D).

As GTCS in mice usually reflect involvement of the limbic system, including the hippocampus, we sought to additionally characterize the epilepsy phenotype of *Hcn1^G380D/+^* and *Hcn1^M142I/+^* mice by performing morphological analysis and immunohistochemical labeling for known hippocampal markers of epilepsy. In line with our video-ECoG findings, we observed upregulation of neuropeptide Y in dentate gyrus mossy fibers, alterations in granule cell layer morphology and extracellular matrix deposition in the dentate gyrus, as well as increased hippocampal gliosis in a majority of mutant animals from both lines (Figure 3 – Figure supplement 1). Such findings are consistent with the presence of spontaneous seizures and altered excitability within the hippocampal circuit (Houser, 1990; Marksteiner et al., 1990; Stringer, 1996; Magagna-Poveda et al., 2017). While the site of seizure onset may differ between species, with a more prevalent role of neocortex in human compared to mouse, these results confirm that *Hcn1^G380D/+^* and *Hcn1^M142I/+^* mice are an overall appropriate model for human *HCN1*-associated DEE, and that the two mutations are causal to the epilepsy phenotype.

### Effects of mutations on HCN1 expression and the intrinsic properties of hippocampal CA1 pyramidal neurons

Prior *in vitro* studies of the human G391D and M153I variants using heterologous expression systems have demonstrated the functional impact of these mutations on HCN1 channels (Marini et al., 2018). HCN1 subunits with the G391D mutation fail to express current on their own, but when mixed with WT subunits (as occurs in patients and heterozygote *Hcn1^G380D/+^* mice) yield, in the ∼20% fraction of cells that show measurable currents, a “leaky” channel with significantly decreased current density but a greatly increased voltage- and time-independent inward, depolarizing current component. In contrast, the M153I mutation only slightly reduces current density, but accelerates the opening kinetics and shifts the voltage dependence of the HCN1 channel to more depolarized potentials (midpoint of activation is shifted by +36 mV for homomeric mutant channels, and by +12 mV for heteromeric M153I/WT channels). However, it is not known how these mutations affect HCN1 expression and neuronal physiology in a native environment, which includes both the HCN channel auxiliary subunit TRIP8b and other channel modulators. To address this question, we performed immunohistochemical labeling of HCN1 protein and patch-clamp recordings from CA1 pyramidal neurons in *Hcn1^G380D/+^* and *Hcn1^M142I/+^* mice and compared their properties with WT littermate controls.

HCN1 antibody labeling of the hippocampus in WT animals showed the typical distribution of HCN1 subunits in CA1 pyramidal neurons (Figure 4A), where the channel is present in a gradient of increasing expression along the somatodendritic axis of the apical dendrite (Lörincz et al., 2002). This gradient of expression was preserved in brains from both mutant lines, although labeling revealed a substantial decrease in overall HCN1 protein levels in *Hcn1^G380D/+^* brains, with a smaller but still significant decrease observed in *Hcn1^M142I/+^* brains (Figure 4B). In whole-cell patch-clamp recordings, such results were matched by a substantial reduction in the voltage sag observed in response to hyperpolarizing current injections, a hallmark of *I*_h_ activity in neurons (Figure 4C,D). The voltage sag was absent in *Hcn1^G380D/+^* CA1 pyramidal neurons and significantly reduced in *Hcn1^M142I/+^*. The lack of voltage sag in *Hcn1^G380D/+^* neurons could be due to decreased HCN1 protein expression, a dominant negative effect of HCN1-G380D subunits on channel function and/or the increased instantaneous component of HCN channels containing HCN1-G380D subunits, which would cause the channel to conduct an inward Na^+^ “leak” current with no voltage dependence (Marini et al., 2018).

**Figure 4.**
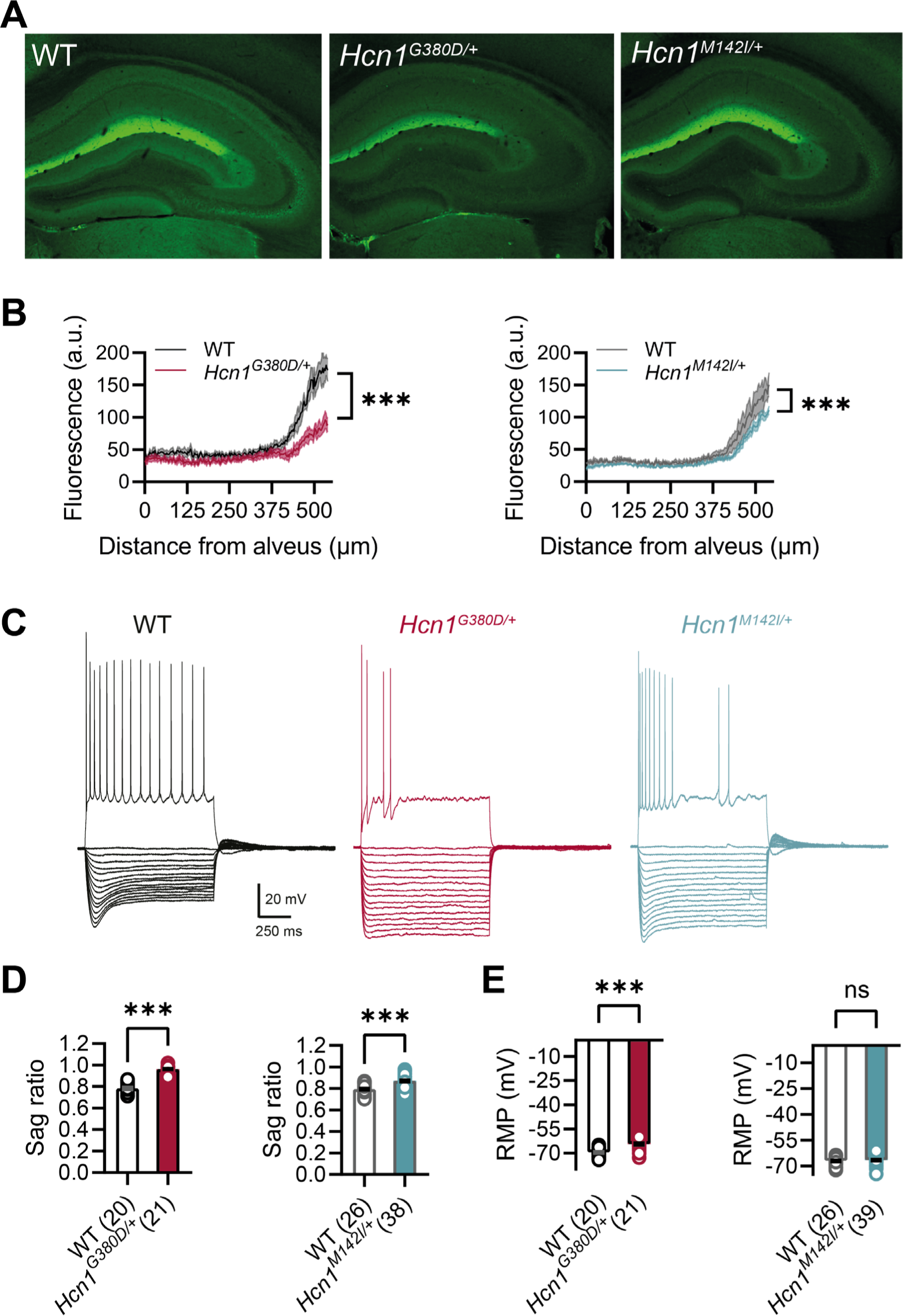
HCN1 protein expression and Ih-dependent voltage sag in CA1 pyramidal neurons are reduced in *Hcn1^G380D/+^* and *Hcn1^M142I/+^* mice. A) Immunofluorescent labeling of HCN1 protein in hippocampus from adult *Hcn1^G380D/+^*, *Hcn1^M142I/+^* and respective WT littermates. B) Quantification of fluorescent signal along the somatodendritic axis of pyramidal neurons demonstrates a significant decrease in protein expression in the distal dendrites, the normal site of greatest HCN1 expression (*** *P* < 0.001, effect of *genotype* x *distance* from alveus after 2-way RM ANOVA; n = 4-7 animals for each sample, n = 2-4 slices per animal; measurements expressed in arbitrary units, a.u.). C) Sample traces from whole-cell current-clamp recordings in hippocampal CA1 pyramidal neurons from WT, *Hcn1^G380D/+^* and *Hcn1^M142I/+^* animals. Holding potential was –70 mV for all samples, with a series of hyperpolarizing and depolarizing current steps in 25 pA increments applied in the negative and positive direction (only depolarizing step at +200 pA is shown). D) Voltage sag in response to hyperpolarizing current steps was significantly reduced in both mutant lines (expressed as sag ratio, see Methods; G380D: WT = 0.78 ± 0.01 *vs. Hcn1^G380D/+^* = 0.96 ± 0.01; M142I: WT = 0.79 ± 0.01 *vs. Hcn1^M142I/+^* = 0.87 ± 0.01; *** *P* < 0.001, Student’s *t* test). E) Resting membrane potential (RMP) was shifted to significantly more positive values in neurons from *Hcn1^G380D/+^* but not *Hcn1^M142I/+^* animals, compared to WT littermates (G380D: WT = -69.55 ± 0.55 mV *vs. Hcn1^G380D/+^* = -64.43 ± 0.71 mV; M142I: WT = -67.08 ± 0.50 mV *vs. Hcn1^M142I/+^* = -66.59 ± 0.51 mV, *** *P* < 0.001, Student’s *t* test; (D-E) number of cells is indicated in the bar graphs; number of animals used: G380D WT n = 6, *Hcn1^G380D/+^* n = 5; M142I WT n = 6, *Hcn1^M142I/+^* n = 5). Data represent mean ± S.E.M.

As wild-type HCN1 channels are partially open at the resting membrane potential (RMP) of most cells, including CA1 pyramidal neurons, they exert a well-characterized action to shift the RMP to more positive potentials and decrease the input resistance. Surprisingly, despite the loss of HCN1 protein expression, CA1 pyramidal neurons of *Hcn1^G380D/+^* mice displayed a significant *depolarization* in the RMP; in contrast, there was no change in RMP in *Hcn1^M142I/+^* neurons (Figure 4E). The more positive RMP of *Hcn1^G380D/+^* mice may reflect the depolarizing action of the increased voltage-independent “leak” current seen with these channels and/or secondary changes in the function of other neuronal conductances (Bleakley et al., 2021). Input resistance was unaltered in both mutant lines (G380D: WT = 137.8 ± 7.1 MΩ, n = 20 cells, *vs Hcn1^G380D/+^* = 153.7 ± 7.8 MΩ, n = 21; M142I: WT = 128.4 ± 9.5 MΩ, n = 25, *vs Hcn1^M142I/+^* = 125.5 ± 7.9 MΩ, n = 38). For *Hcn1^G380D/+^* mice this may again reflect the offsetting actions of increased “leak” current and decreased channel expression, or alternatively, offsetting effects due to alterations in the expression of other types of channels.

The possibility that the expression of mutant HCN1 channels may cause secondary changes in CA1 pyramidal neuron function is consistent with our finding that CA1 cells in both mouse lines show an impaired ability to fire action potentials, a phenotype not observed with acute application of the HCN channel specific blocker ZD7288 (Gasparini and DiFrancesco, 1997; and Figure 4 – Figure supplement 1). Thus, application of positive current steps from a common holding potential of –70 mV revealed an abnormal firing pattern in CA1 neurons from both lines (Figure 4C), wherein a majority of *Hcn1^G380D/+^* neurons and a significant fraction of *Hcn1^M142I/+^* neurons showed a reduction in the number of action potentials fired during one-second-long depolarizing current steps (fraction of neurons with abnormal firing pattern: 76% *Hcn1^G380D/+^ vs* 37% *Hcn1^M142I/+^ vs* 2% in the combined set of WT littermate neurons). In contrast, rheobase was unaltered in both lines, when measured from a holding potential of –70 mV (G380D: WT = 100 ± 9.6 pA, n = 20, *vs Hcn1^G380D/+^* = 114.3 ± 8.0 pA, n = 21; M142I: WT = 101.0 ± 6.8 pA, n = 25 *vs Hcn1^M142I/+^* = 98.6 ± 6.4 pA, n = 38). No differences were observed between males and females for any of the measures assessed in either mouse line, hence all reported data were pooled across sexes.

Overall, the observed changes reflect subtle alterations in the intrinsic properties of pyramidal neurons, which do not immediately explain the hyperexcitability and epilepsy phenotype observed in either mutant mouse line. The reduced ability of *Hcn1^G380D/+^* pyramidal neurons to maintain sustained firing may, in fact, act to limit circuit hyperexcitability in these animals – as reflected in the lower average maximum seizure grade reached by *Hcn1^G380D/+^* mutants both during spontaneous and drug-induced seizure events (Figure 3B,D; and see Figure 7C below), compared to *Hcn1^M142I/+^* animals. These results suggest that altered activity in non-pyramidal neurons may also play a role in the epilepsy phenotype of the mutants.

### Severe impairment in the axonal localization of HCN1 protein in Hcn1^G380D/+^ PV+ interneurons

As HCN1 subunits are highly expressed in PV+ interneurons in the brain, in addition to their expression in pyramidal neurons, we examined whether this expression is altered by the seizure-causing mutations. Whereas the highest site of expression of HCN1 subunits is at the distal portion of apical dendrites of pyramidal neurons, as detailed above, HCN1 is most strongly enriched in the axonal terminals of PV+ interneurons. The exquisite targeting of HCN1 protein to axonal terminals in PV+ interneurons can be particularly appreciated in basket cells from the cerebellar cortex (Figure 5). Here, HCN1 protein accumulates in a specialized structure, named the “pinceau”, representing the axonal termination of basket cells onto the axon initial segment of Purkinje neurons. Surprisingly, at this location, we found a complete loss of HCN1 protein staining in *Hcn1^G380D/+^* animals, while the distribution in *Hcn1^M142I/+^* appeared relatively normal (Figure 5, top row). This result is unexpected, as levels of HCN1 protein expressed from the WT allele should be normal, providing at least a half dosage in *Hcn1^G380D/+^* cells, and suggests that the mutation may exert a dominant negative effect by coassembling with WT HCN1 subunits, resulting in improper trafficking and/or premature degradation.

**Figure 5.**
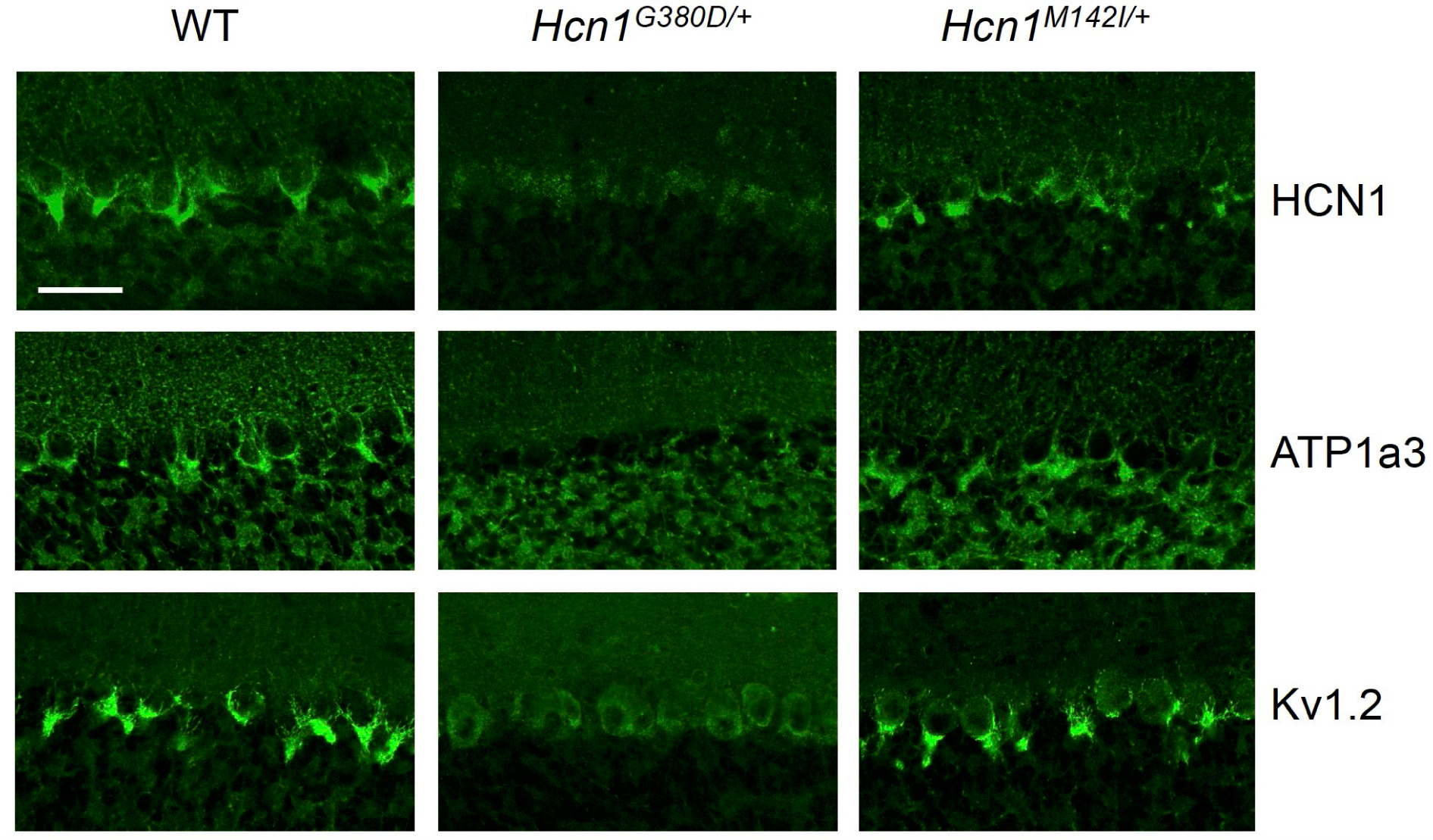
Alterations in HCN1 localization in cerebellar basket cell *pinceau* with accompanying loss of Kv1.2 and ATP1a3 in *Hcn1^G380D/+^* but not *Hcn1^M142I/+^* mice. Immunofluorescent labeling of coronal hindbrain sections from adult mouse brain. The cerebellar cortex of WT, *Hcn1^G380D/+^* and *Hcn1^M142I/+^* mice is shown. Top row, HCN1 protein labeling; middle row, Na^+^/K^+^ pump labeling (protein isoform ATP1a3); bottom row, Kv1.2 protein labeling. The PV+ IN (basket cell) terminal *pinceau* surrounding the Purkinje cell axon initial segment is labeled by HCN1, ATP1a3 and Kv1.2 antibodies in WT and *Hcn1^M142I/+^*, while no labeling is present in *Hcn1^G380D/+^* brains. Scale bar = 50 μm in all panels.

Accumulation of misfolded protein in the endoplasmic reticulum or Golgi apparatus may also impair the trafficking of proteins other than HCN1. As several other proteins and channels are known to specifically localize to the pinceau, we used labeling for one such channel (i.e., K_v_1.2) to assess whether the normal pinceau channel architecture is disrupted in *Hcn1^G380D/+^* brains. As shown in Figure 5 (bottom row), K_v_1.2 labeling of the pinceau was also completely lost, similar to HCN1 protein labeling, in *Hcn1^G380D/+^* but not *Hcn1^M142I/+^* brains. The downregulation of K_v_1.2 does not appear to be due to co-regulation with HCN1, as normal K_v_1.2 labeling is still present in the cerebellum of HCN1 knockout mice (Figure 5 – Figure supplement 1) and the expression of the two channels at the cerebellar pinceau was shown to be independently regulated (Kole et al., 2015). As HCN1 channels in PV+ axonal terminals are thought to functionally interact with the Na^+^/K^+^ pump (Roth and Hu, 2020), we further sought to test for the presence of ATP1a3 expression at the pinceau. As shown in Figure 5 (middle row), similar to HCN1 and K_v_1.2, there was a complete loss of ATP1a3 labeling in *Hcn1^G380D/+^* but not *Hcn1^M142I/+^* animals, compared to WT controls.

The profound disruption of normal pinceau architecture in cerebellar basket cells suggests that, in the case of HCN1-G380D, misfolding of the mutated protein may be causing cellular stress (presumably endoplasmic reticulum stress, due to impaired trafficking and/or increased protein degradation). Such toxicity effects may explain the higher severity of the *Hcn1^G380D/+^* phenotype, which includes altered anatomy and behaviors beyond the occurrence of seizures and epilepsy.

Similar to what was observed in the apical dendrites of CA1 pyramidal neurons (Figure 4A,B), immunolabeling of HCN1 protein in the axon terminals of hippocampal PV+ interneurons was strongly reduced in *Hcn1^G380D/+^* animals, although not completely absent (Figure 6A). Since endogenous protein labeling is unable to distinguish between WT and mutant subunits, we sought to further probe for the trafficking of mutant HCN1 subunits in hippocampal PV+ interneurons using a viral transduction approach. To this end, we used mice that express Cre recombinase specifically in PV+ interneurons (*Pvalb-Cre* line) and stereotaxically injected into the hippocampal CA1 area an adeno-associated virus (AAV) driving Cre-dependent expression of hemagglutinin (HA)-tagged HCN1 subunits (AAV-DIO-HA-HCN1). As shown in Figure 6B, following injection of either AAV-DIO-HA-HCN1(WT) or AAV-DIO-HA-HCN1(M142I), staining with anti-HA antibodies revealed efficient targeting and expression of virus-driven protein in the axonal terminals of PV+ interneurons. In stark contrast, injection of AAV-DIO-HA- HCN1(G380D) did not result in detectable levels of HA-tagged protein in axonal terminals, while protein accumulation was observed in the soma of targeted neurons (Figure 6B). These results further underscore the adverse effect of the latter variant on the biosynthesis of the HCN1 protein, and suggest that disruption of PV+ interneuron function, due to HCN1 protein misfolding or altered trafficking, may represent an important component of neuronal circuit dysfunction in HCN1-linked epilepsy.

**Figure 6.**
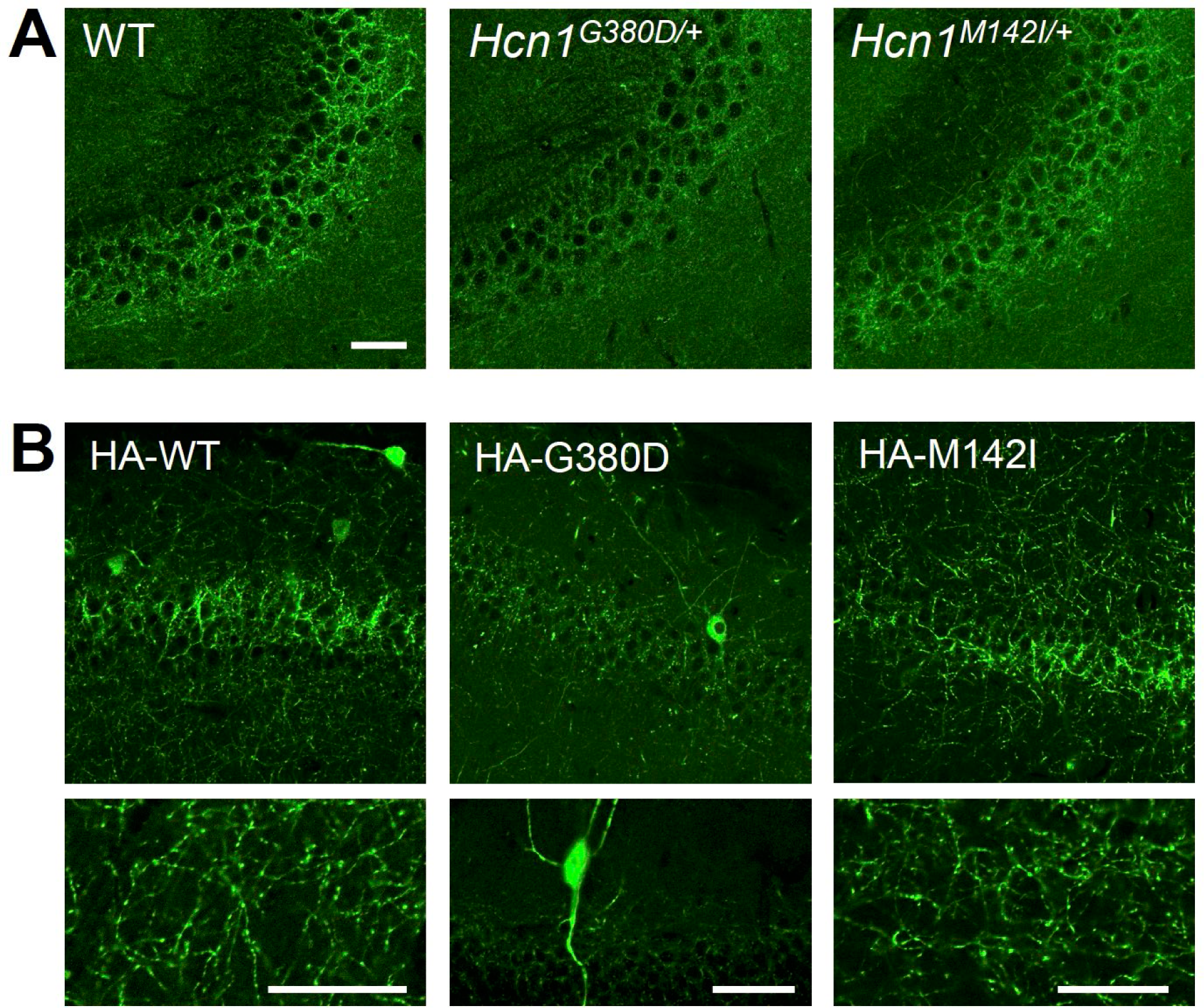
Viral targeting of HCN1 subunits to PV+ interneuron terminals in hippocampus shows impaired trafficking of G380D mutant protein Immunofluorescent staining of midcoronal sections from adult mouse brain. A) Labeling of HCN1 protein in the hippocampus of *Hcn1^+/+^* (WT), *Hcn1^G380D/+^* and *Hcn1^M142I/+^* mice. Images show a close-up of the pyramidal cell layer of area CA3, with the PV+ interneuron axonal terminals visible. B) Anti-HA tag labeling of virally expressed HCN1 protein after stereotaxic injection into hippocampal area CA1 of adult Pvalb-Cre mice. Panels show distribution of HA- HCN1-WT (HA-WT), HA-HCN1-G380D (HA-G380D) and HA-HCN1-M142I (HA-M142I) protein at 21 days after injection. Scale bars = 50 μm (same bar for all panels in A and upper panels in B; lower panels in B show individual bars for higher magnification images). Note the accumulation of HA-HCN1-G380D protein in the soma and dendrites of PV+ interneurons, with minimal or absent targeting to axon terminals.

### Paradoxical response of Hcn1^G380D/+^ and Hcn1^M142I/+^ mice to antiseizure medications

Clinical reports from at least four different patients with *HCN1*-linked DEE (carrying mutations M305L, G391D, and I380F; Marini et al., 2018; Bleakley et al., 2021; C. Marini, personal communication, March 2021) consistently revealed worsening of seizures after administration of either lamotrigine or phenytoin. A particularly striking example is provided by the G391D mutation, identified in two unrelated patients with similar course of disease. Both G391D patients had neonatal seizure onset, characterized by daily asymmetric tonic seizures with apnea and cyanosis, severe developmental delay, and died between 14-15 months of age. Both patients also showed a paradoxical response to phenytoin, which caused the induction of status epilepticus. Given this precedent, we hypothesized there may be systematic similarities between the adverse pharmacological response profile of HCN1 patients and *Hcn1^G380D/+^* and *Hcn1^M142I/+^* mutant mice, which would further endorse their use as models with construct, face and potentially predictive validity.

Consistent with our hypothesis, we found that administration of lamotrigine (23 mg/kg, i.p.) induced convulsive seizures, defined as grade 3 or higher on a modified Racine scale (see Materials and Methods), within 90 min from injection in 11/15 *HCN1^G380D/+^* animals (Figure 7A,C, left). Several animals had multiple seizure bouts, ranging from rearing with forelimb clonus to wild running and jumping seizures, during the period of observation. Video-ECoG recordings were performed on a subset of animals (2/15 mutant and 2 WT control animals) for a total of 24 h after drug injection, confirming the presence of GTCS-like activity on ECoG in both mutants (Figure 7A) but no seizure activity in control littermates (data not shown). Similar outcomes were seen in *Hcn1^M142I/+^* animals, where convulsive seizures were observed in 6/9 mutant mice in response to lamotrigine (Figure 7A,C, right). Of note, four of the six *Hcn1^M142I/+^* animals with adverse response had grade 6 seizures, again higher than the maximum seizure grade observed in *Hcn1^G380D/+^* mice. (Figure 7C). Video-ECoG recordings performed on three *Hcn1^M142I/+^* mutants confirmed the presence of GTCS electrographic activity in 2/3 animals (Figure 7A). In both lines, all seizure events recorded through video-ECoG were reflected in behavioral manifestations, implying that data collected through observation in non-implanted animals are likely to have captured all seizure events generated in response to lamotrigine injection.

**Figure 7:**
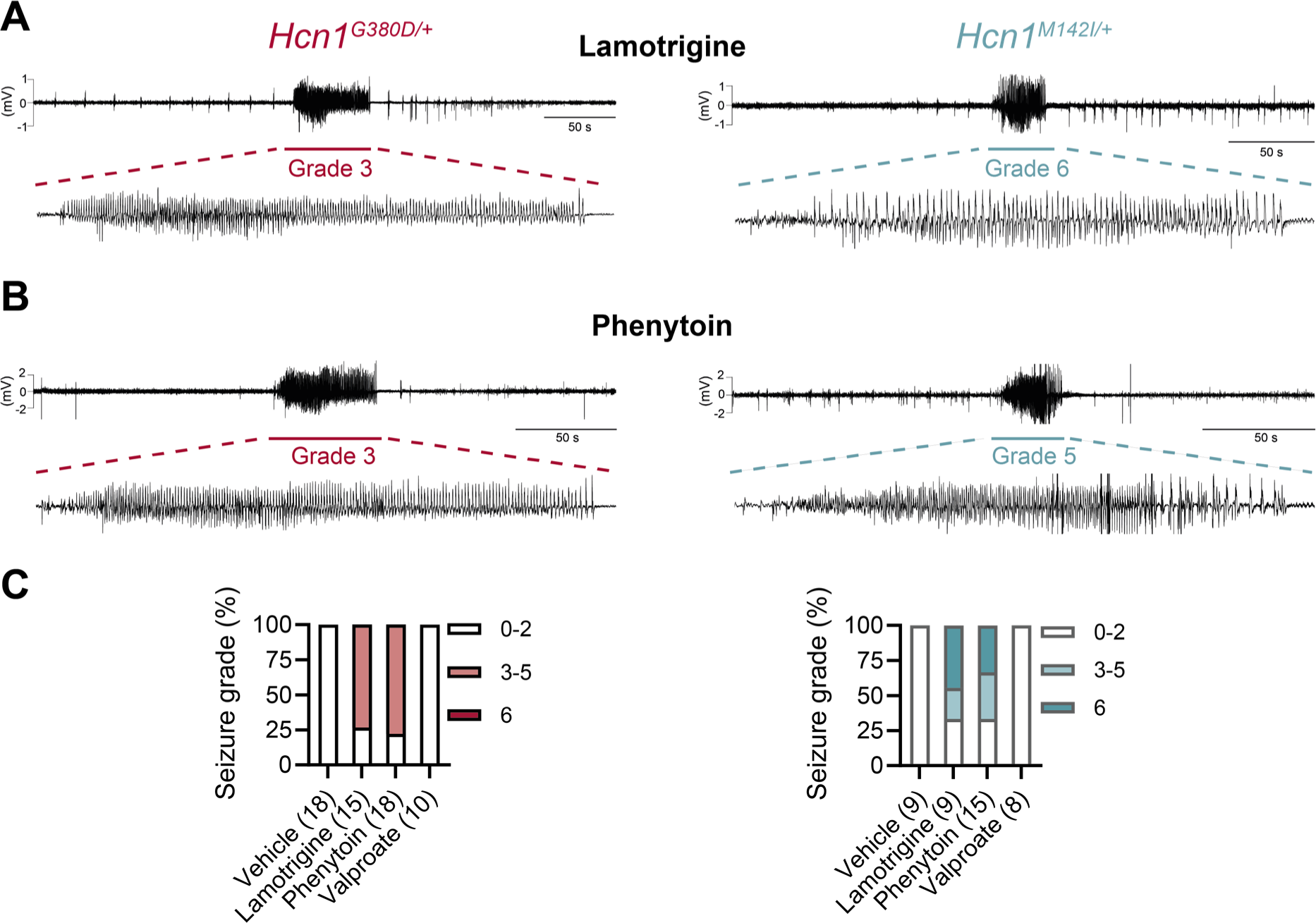
Anticonvulsant-drug induced seizures in *Hcn1^G380D/+^* and *Hcn1^M142I/+^* mice. A) Example ECoG trace for lamotrigine-induced grade 3 seizure in *Hcn1^G380D/+^* (left) and grade 6 seizure in *Hcn1^M142I/+^* mice (right), with seizure shown at expanded time scale below. B) Phenytoin-induced grade 3 seizure in *Hcn1^G380D/+^* (left) and grade 5 seizure in *Hcn1^M142I/+^* mice (right). C) Percentage of *Hcn1^G380D/+^* (left) and *Hcn1^M142I/+^* (right) animals experiencing convulsive seizures (grade 3 and higher) upon administration of vehicle, lamotrigine, phenytoin, or sodium valproate, with highest seizure grade indicated by color. As in the case of spontaneous seizures, *Hcn1^M142I/+^* animals showed an overall higher seizure severity in response to drug administration. No convulsive seizures were observed in response to vehicle or sodium valproate during the observation period of the experiment (see Materials and Methods). Total number of animals tested for each condition is reported in parentheses (see Results for number of video-ECoG recordings *vs* behavioral observations).

Results with phenytoin (30 mg/kg PE, i.p.) were even more striking, as 14/18 mutant animals tested from the *Hcn1^G380D/+^* line developed convulsive seizures within 2.5 h of injection (this number includes 11/14 animals with no prior drug exposure and 3/4 animals previously exposed to lamotrigine; Figure 7B,C, left). Latency to first seizure after injection was longer than observed with lamotrigine, in line with the slower time to peak plasma concentration for phenytoin. In addition, multiple seizures were observed during a period up to six hours after injection, again consistent with the longer half-life of phenytoin compared to lamotrigine (Markowitz et al., 2010; Hawkins et al., 2017).

Among *Hcn1^M142I/+^* animals, we observed seizures in 10/15 mice following phenytoin injection, with similar temporal pattern and ECoG signatures compared to *Hcn1^G380D/+^* mice (this number includes 6/8 animals without prior drug exposure, and 4/7 animals previously exposed to lamotrigine, Figure 7B,C, right).

All the *Hcn1^G380D/+^* or *Hcn1^M142I/+^* animals tested, either in response to lamotrigine or phenytoin, were randomized to receive a control vehicle injection either one week before or one week after the drug administration session. None had convulsive seizures in response to vehicle injection during the period of observation (4 h for behavioral experiments, and 24 h for video-ECoG experiments), further confirming that the seizures observed in the period following drug administration were a direct consequence of the pharmacological challenge (Figure 7C). As an additional control, we tested the effects of a third anticonvulsant, namely sodium valproate (VPA), which has not been reported to worsen seizures in any of the HCN1 patients in our database. Administration of sodium valproate (250 mg/kg, i.p.) did not result in the occurrence of convulsive seizures in any of the *Hcn1^G380D/+^* or *Hcn1^M142I/+^* animals tested (Figure 7C).

However, use of 250 mg/kg VPA resulted in profound sedation of *Hcn1^G380D/+^* mice, which was not seen in *Hcn1^M142I/+^* mice or WT littermate controls, with 9/10 *Hcn1^G380D/+^* animals failing to respond to gentle touch or passive moving of the tail 45 min after drug injection *vs* 2/8 *Hcn1^M142I/+^* and 0/8 WT animals (note that tail pinching still elicited a response in all mice tested).

What may be the mechanism by which lamotrigine and phenytoin induce seizures in HCN1 mutant mice? Although both drugs act as Na^+^ channel blockers, lamotrigine is also widely considered an HCN channel activator, a property thought to contribute to its antiepileptic effects (Poolos et al., 2002). Despite multiple reports on its action on *I*_h_ in neurons, showing somewhat inconsistent outcomes (Peng et al., 2010; Huang et al., 2016), no studies have been conducted to directly test the effect of lamotrigine on isolated HCN channels expressed in heterologous systems. To obtain further clarification on the mechanism of action of lamotrigine, we tested the drug on wildtype HCN1 and HCN2 channels after transient transfection in HEK293T cells. Surprisingly, we found that lamotrigine had no effects on either the midpoint voltage of activation or maximal tail current density of either HCN1 or HCN2 channels. Lamotrigine also had no effect when HCN1 was coexpressed with its auxiliary subunit TRIP8b (Figure 8). In contrast, under the same conditions, lamotrigine exerted its expected inhibitory effect on Na_v_1.5 sodium channel currents (Figure 8 – Figure supplement 1; Qiao et al., 2014). These results clearly show that lamotrigine is not a direct activator of HCN channels, and suggest that any effects reported in neurons are the result of indirect regulation of *I*_h_, perhaps secondary to Na^+^ channel block, and likely contingent on cell-specific, state-dependent signal transduction pathways. Thus, it is more likely the effects of lamotrigine on *Hcn1^G380D/+^* and *Hcn1 ^M142I/+^* animals are due to its action to directly block Na^+^ channels – similar to the effects of phenytoin – rather than an enhancement of HCN currents.

**Figure 8:**
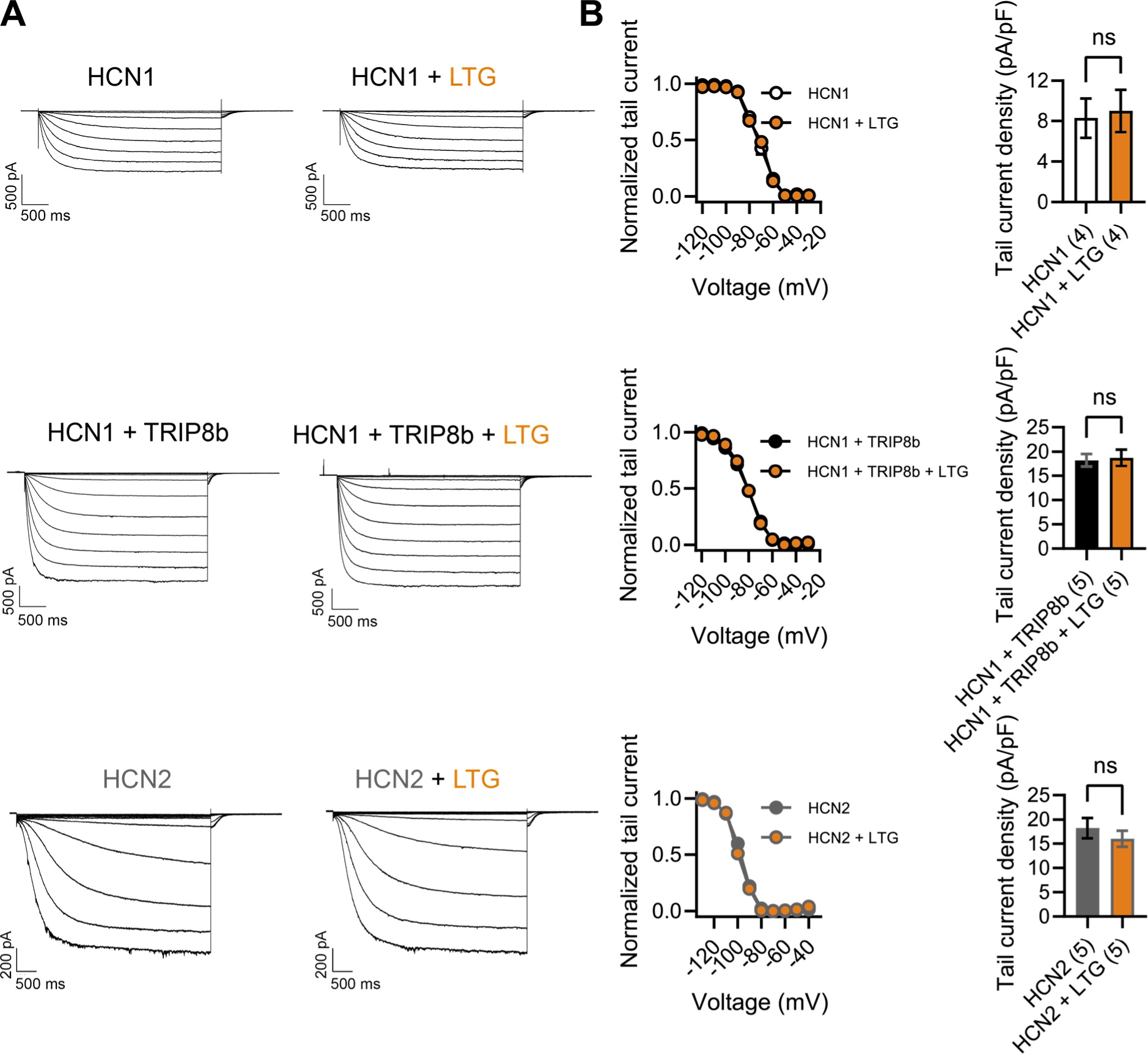
Lamotrigine has no direct effect on HCN1 or HCN2 channel activity. A) Sample current traces from whole cell voltage-clamp recordings in HEK293T cells transiently expressing HCN1 (top), HCN1 and TRIP8b (middle) or HCN2 (bottom), in the absence or presence of bath applied lamotrigine (LTG, 100 μM). Voltage step protocol for HCN1 and HCN1+TRIP8b: holding potential was -20 mV (1 s), with steps from -30 mV to -120 mV (-10 mV increments, 3.5 s) and tail currents recorded at -40 mV (3.5 s); for HCN2: holding potential was similarly −20 mV (1 s), but steps were applied from −40 mV to −130 mV in -15 mV increments (5 s) and tail currents recorded at −40 mV (5 s). B) Tail current activation curves (left) and tail current density (right) group data obtained for each condition. Lines show data fitting with a Boltzmann function (see Material and Methods). Data show that neither the midpoint voltage of activation (V1/2) nor the maximum tail current density was altered by LTG addition. (ns = not significant, Student’s *t* test; see Figure 8 – source data 1 for numerical values) Data represent mean ± S.E.M.

While we have at present no available indication that either of the two M153I patients showed adverse effects to lamotrigine or phenytoin treatment, our results in mice suggest that despite the clear differences in their phenotypes, there may be some fundamental similarities in the disease mechanisms at play both in patients and mouse models for *HCN1*-linked DEE.

### Reduced efficacy of HCN/I_h_-blocking compounds on HCN1-G391D channels and Hcn1^G380D/+^ neurons

The observation that several of the known *HCN1* variants, when tested for their properties *in vitro* (Nava et al., 2014; Marini et al., 2018) or *in vivo* (Bleakley et al., 2021), exhibit an increase in “leak” current suggests that use of HCN channel blockers may be beneficial in the treatment of certain *HCN1* patients. Several such blockers exist, although at present none is able to efficiently cross the blood brain barrier. More importantly, all currently available inhibitors that are specific for HCN channels, such as ZD7288, ivabradine, zatebradine, cilobradine and their derivatives (Melchiorre et al., 2010; Del Lungo et al., 2012), are thought to act through the same mechanism, namely as pore blockers by interacting with residues located within the pore cavity. At the same time, many of the most severe *HCN1* variants, including G391D, are located in the S6 transmembrane domain of the channel, which forms the inner lining of the pore. Such observations raise the question whether pathogenic mutations in S6 may decrease the efficacy of HCN pore-blocking compounds.

The structural models presented in Figure 9F illustrate the proximity of the G391D mutation to the known binding site of ZD7288 in the HCN pore cavity, wherein the residue immediately preceding G391 (V390 in human HCN1, corresponding to V379 in mouse) was shown to critically affect the sensitivity of the channel to the drug (Cheng et al., 2007). The presence of charged aspartate residues in G391D-containing subunits leads to a disruption in the symmetry of the pore in heteromeric channels containing at least one WT subunit, and consequently in the relative distance between the V390 side chains facing the cavity. To test whether this predicted disruption in the ZD7288 binding site leads to a reduced efficacy of the drug, we expressed heteromeric HCN1 channels containing WT and G391D mutant subunits in HEK cells. As illustrated in Figure 9A,B, compared to the effect of ZD7288 to reduce the current expressed by WT HCN1 channels by ∼67%, the drug was minimally effective in blocking either the instantaneous or the time-dependent current component generated by mutant WT/G391D channels (∼15% inhibition). Addition of Cs^+^, another HCN channel blocker (albeit a less specific one, compared to ZD7288) with a different mechanism of action and binding site (DiFrancesco, 1982), led to a reduction in the current expressed by both WT and mutant channels, confirming that the observed currents were indeed generated by HCN1 WT/G391D heteromeric channels.

**Figure 9:**
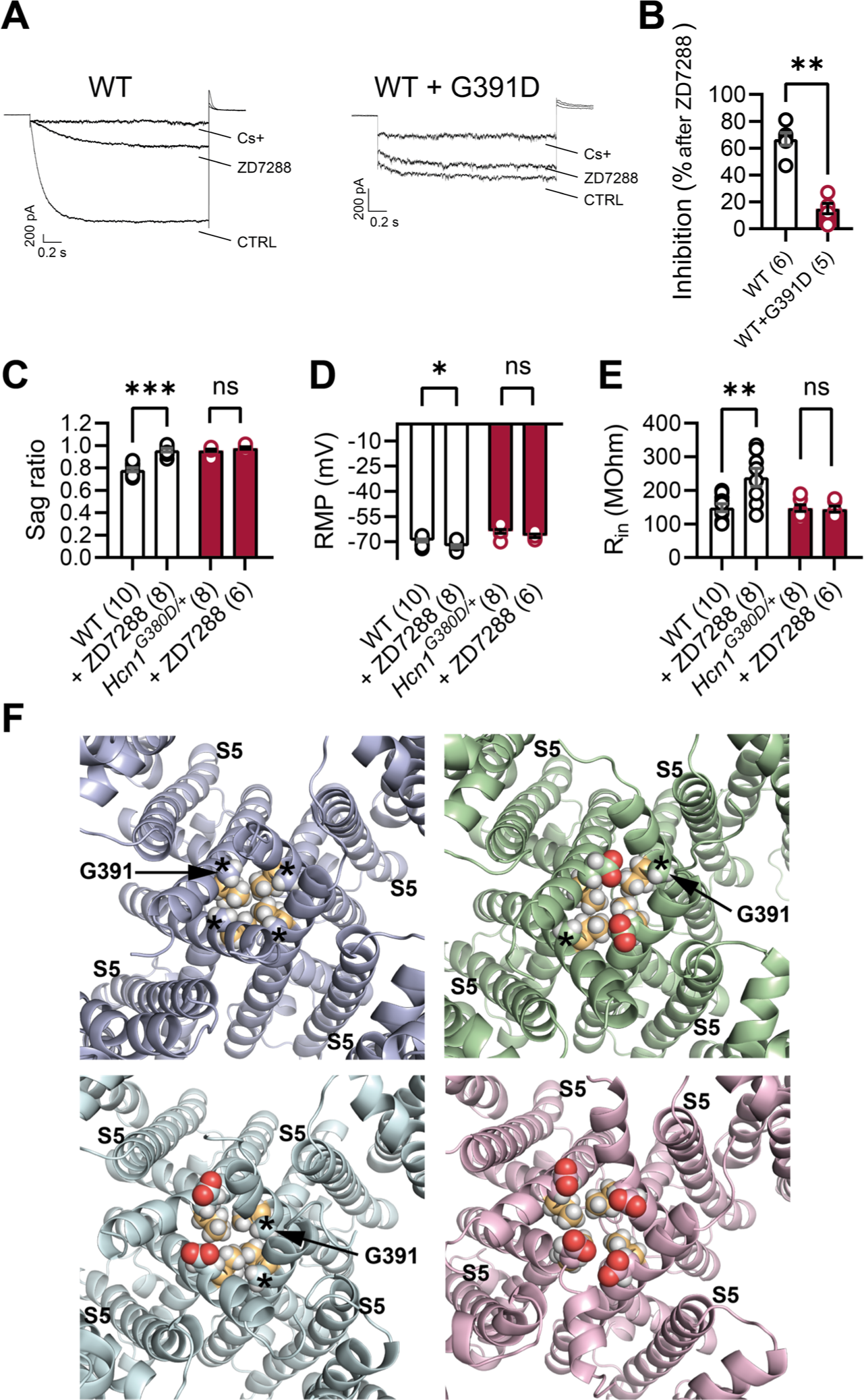
Lack of effect of ZD7288 on *Hcn1^G380D/+^* neurons and heterologously expressed hHCN1-WT/G391D channels. A) Whole cell voltage clamp recordings in HEK293T cells heterologously expressing either human HCN1 (hHCN1) WT or heteromeric hHCN1 WT/G391D channels. Sample current traces are shown, in response to a voltage step to -110 mV from a holding potential of -20 mV, before (control, CTRL) and after full block from ZD7288 (30 μM) or CsCl (5 mM) bath application. B) Bar plot with % inhibition from group data (WT= 66.83 ± 4.5% *vs* WT/G391D = 15.00 ± 3.89 %; *** *P* < 0.001, Mann Whitney *U* test). Data represent mean ± S.E.M. C-E) The HCN blocker ZD7288 (10 μM) abolished sag (C) significantly hyperpolarized RMP (D) and increased input resistance (E) in WT, but did not alter any of these parameters *Hcn1^G380D/+^* neurons. (** *P* < 0.01, *** *P* < 0.001, paired *t* test; number of recorded cells is indicated in the bar graphs; number of animals used: WT n = 4, WT+ZD7288 n = 3, *Hcn1^G380D/+^* n = 3, *Hcn1^G380D/+^*+ZD7288 n = 3; see Figure 9 – source data 1 for numerical values) F) Snapshots of the human HCN1 pore region structure at the end of a 100 ns Molecular Dynamics (MD) simulation run, with or without introduction of the G391D mutation in two or four subunits. Top left, WT; top right, heteromer with G391D variant on two opposite subunits; bottom left, heteromer with G391D variant on two adjacent subunits; bottom right, homomer with G391D variant on all four subunits. The outer pore helix S5 is labeled (S5), with asterisks indicating the position of the G391 residue in all cases. Carbonyls of modeled G391D residues are colored in red. Side chains of V390 residues are colored in light orange.

Parallel experiments in brain slices, where bath application of ZD7288 is the most common method used to isolate the contribution of *I*_h_ to neuronal physiology, demonstrate the additional difficulties posed by the limited efficacy of the drug in the mutant. As shown in Figure 9C-E, application of the compound did not modify either the RMP or input resistance in *Hcn1^G380D/+^* neurons, while appropriately eliminating voltage sag, hyperpolarizing the RMP, and increasing input resistance in WT neurons. These results leave unanswered the question whether the positive shift in RMP in *Hcn1^G380D/+^* neurons is directly due to “leaky” HCN1 channels or is a secondary response to changes in other conductances, further underscoring the need for a radically different approach in dealing both with the therapeutic treatment and experimental investigation of *HCN1*- linked DEEs.

## Discussion

The speed and precision of next generation sequencing (NGS) allows the efficient genetic assessment of children that present with early onset epilepsies. The overarching goal of such precision medicine is to identify the specific molecular and network mechanisms that link distinct epilepsy mutations in individual genes to particular disease profiles, resulting in more effective gene- or mutation-specific therapeutic strategies. Despite the fact that dysfunction in ion channels carrying missense mutations can be promptly modeled in heterologous expression systems, such assessments do not sufficiently reflect the dysfunction caused by the mutations *in vivo*, preventing the direct translation of such datasets into applicable clinical strategies. In our study, we set out to fill this information gap by designing new preclinical models, which reproduce key disease features of *HCN1-*linked DEE. The *Hcn1^G380D/+^* and *Hcn1^M142I/+^* knock-in mice we generated display not only an appropriate and robust epilepsy phenotype, along with comorbid behavioral abnormalities, but illustrate the difficulties intrinsic to the treatment of genetic developmental epilepsies, including the adverse response of individuals with *HCN1* mutations to conventional antiseizure medications, and the potential for altered drug sensitivity of mutant channels. These results underscore how mutations in voltage-gated ion channels can fundamentally alter brain development and the response of neuronal circuits to pharmacological challenge. In this scenario, our models provide a useful system for testing mechanistic hypotheses and developing new therapeutic approaches.

A notable feature of our models and patients with *HCN1* mutations is the paradoxical induction of seizures by lamotrigine and phenytoin. A similar phenomenon has been reported both in Dravet syndrome patients carrying *SCN1A* mutations (Perucca and Perucca, 2019) and in mice modeling *Scn1A* haploinsufficiency (*Scn1A^+/-^*; Hawkins et al., 2017). In Dravet syndrome, loss-of-function mutations in the *SCN1A* sodium channel gene are thought to promote seizures by exerting a net effect to reduce the excitability of inhibitory interneurons (Catterall, 2018). This has led to the suggestion that certain anticonvulsant drugs acting as Na^+^ channel blockers paradoxically exacerbate seizures in this syndrome as a result of a synergistic action to further suppress inhibition. Interestingly, the axon-specific expression of HCN1 channels in PV+ interneurons matches the similarly polarized subcellular distribution of voltage-gated Na^+^ channels, wherein Na_v_1.1 (encoded by *SCN1A*) is the predominant subtype expressed in the axon of PV+ interneurons (Ogiwara et al., 2007; Dutton et al., 2013; Ogiwara et al., 2013; Hedrich et al., 2014; Hu and Jonas, 2014). To maintain ionic homeostasis during repetitive firing, PV+ interneuron axons also strongly express Na^+^/K^+^-ATPases (Peng et al., 1997), which will activate in response to Na^+^ entry leading to membrane hyperpolarization due to electrogenic pump activity. Recent pharmacological studies *in vitro* have proposed that axonal expression of HCN channels can counteract the hyperpolarizing Na^+^ pump current during repetitive firing, allowing for fast action potential firing and propagation (Roth and Hu, 2020). Thus, blockade of HCN channel activity in PV+ interneuron axon terminals results in a failure in action potential propagation and reduced perisomatic inhibition onto target excitatory neurons (Southan et al., 2000; Aponte et al., 2006; Roth and Hu, 2020). Since the only HCN subtype expressed in PV+ interneuron terminals is HCN1, diminished GABAergic input onto excitatory neurons as a result of HCN1 protein dysfunction could contribute to excitation/inhibition imbalance and epilepsy. Such an effect could be exacerbated by further inhibition of Na^+^ channel activity by antiseizure medications, similar to what is thought to occur in Dravet syndrome. HCN1 subunits also substantially contribute to regulating the excitability of a second class of inhibitory neurons, namely somatostatin-positive interneurons, which target the dendrites of pyramidal cells (Matt et al., 2011). Here, HCN1 acts along with HCN2 to set the somatic resting potential and modulate the spontaneous activity of this class of interneurons. Future studies aimed at investigating the physiological effects of HCN1 mutations in identified interneuron populations, along with conditional knock-in mouse lines which limit allele expression to select classes of inhibitory neurons, may provide a more definitive answer to the question of how different variants may affect perisomatic and/or dendritic inhibition in *HCN1*-linked epilepsy.

When tested in heterologous expression systems, epilepsy-linked HCN1 mutations have been shown to affect the channel in several different ways. These actions include loss-of-function due to decreased HCN1 channel activation, decreased protein levels, and/or impaired targeting to the plasma membrane. Some of the variants exert a dominant negative effect when a mutant subunit assembles into a heteromeric channel with WT subunits. Conversely, gain-of-function effects include accelerated activation kinetics, depolarized midpoint voltage of activation (as in M153I), and an increased voltage-independent component of the HCN current, i.e., generation of a depolarizing “leak” current (as in G391D). Why do *HCN1* mutations that in heterologous expression systems have divergent effects on channel function equally result in epilepsy, and a similar paradoxical response to certain Na^+^ channel blockers? Our results in *Hcn1^G380D/+^* and *Hcn1^M142I/+^* mice show that both these mutations ultimately led to a net decrease in *I*_h_-dependent voltage sag, and a significant decrease in protein expression levels (Figure 4), despite the M153I variant yielding an apparent gain-of-function channel upon heterologous expression (Marini et al., 2018). This suggests that misfolding and impaired biosynthesis of mutant HCN1 subunits may be a prevalent consequence likely to affect the function of neurons in multiple ways. An increase in cellular stress, for example, would be expected to affect PV+ neurons in particular as they are known to be extremely sensitive to metabolic and oxidative stress (Kann, 2016). Such a mechanism may explain why two variants with seemingly divergent biophysical alterations in channel functional properties ultimately cause overlapping phenotypes.

The divergent characteristics of the two variants do, however, lead to discernible phenotypes, as shown by the overall greater severity of the *Hcn1^G380D/+^* DEE phenotype, and the remarkable sex-dependence of mortality in *Hcn1^M142I/+^* mice (Figure 1). This suggests that, while some fundamental mechanisms are affected in similar ways, there are distinctions in the manner in which developmental trajectories are perturbed by different *HCN1* mutations. As for other known ion channel linked syndromes, such distinctions may explain the variety of disease phenotypes observed in patients carrying *HCN1* pathogenic variants, which range from epileptic encephalopathies to genetic generalized epilepsies and genetic epilepsy with febrile seizures plus (Bonzanni et al., 2018; Marini et al., 2018). An analogous picture emerges from the study of a third *HCN1* variant recently modeled in mice (*Hcn1^M294L/+^*, corresponding to human p.Met305Leu; Bleakley et al., 2021). Similar to HCN1-G391D, this variant generates a channel with a significant “leak” or voltage-independent current component, although protein levels in the *Hcn1^M294L/+^* mouse brain are less dramatically affected. The epileptic phenotype of *Hcn1^M294L/+^* mice is indeed considerably milder than either *Hcn1^G380D/+^* or *Hcn1^M142I/+^* animals.

Intriguingly, however, *Hcn1^M294L/+^* mice are also adversely affected by lamotrigine, which caused the induction of seizures, similar to mutant animals from the two lines presented here. From the therapeutic point of view, the observed mechanistic convergence does offer some hope that patients with genetic HCN1 channel dysfunction can be treated following similar principles or strategies across different variants. A larger convergence in treatment strategies, centered on the degree of interneuronal dysfunction across multiple epilepsies, may also be envisioned to emerge in the future.

Finally, what may such strategies look like? Our data once again stress the difficulty of devising targeted therapeutic approaches for ion channel-associated DEE syndromes using existing tools. As stated above, several of the identified epilepsy-associated *HCN1* variants generate channels with gain of function and/or gain of aberrant function properties, implying that HCN channel blockers may be potentially helpful. At the same time, several of the most severe mutations in *HCN1* epilepsy patients are located in the S6 segment, which forms the lining of the pore cavity, or in the immediate vicinity to the channel’s intracellular gate. These include G391D (Figure 9) and two more variants at the same position, G391C and G391S; as well as M379R, I380F, A387S, F389S, I397L, S399P, and D401H (Nava et al., 2014; Lucariello et al., 2016; Marini et al., 2018; Wang et al., 2019). While we do not have complete information available about the effects of known HCN inhibitors on the other variants, results presented here using ZD7288 on G391D-containing HCN1 channels strongly suggest that the mutations may decrease the efficacy of the pore-blocking compounds, reducing their ability to suppress both the time-dependent and time-independent current component generated by the mutant channels (Marini et al., 2018). There is therefore a need to develop new small molecule compounds that target the HCN1 channels at sites outside the pore cavity.

The available 3D structures of HCN1, combined with molecular dynamics simulations, coarse-grained modeling and molecular docking, suggest that there are many opportunities for such a pharmacological-targeting approach to succeed (Lee and MacKinnon, 2017; Gross et al., 2018; Porro et al., 2019). The structures have indeed revealed unique intracellular regulatory modules, including both the HCN domain and C- linker/CNBD, which may allow for the targeting of HCN channels through allosteric modulation of pore gating properties. Furthermore, the recently determined structures of HCN4, the main cardiac isoform of HCN channels, have demonstrated critical differences in the relative arrangement of such regulatory modules (Saponaro et al., 2021), which may be exploited towards the development of isoform-specific drugs. Compound library screenings for small molecules with HCN1-subtype specificity have already yielded some initial and promising results (McClure et al., 2011; Harde et al., 2019). Other approaches include the design of cell-penetrant peptides, which interfere with the interaction between HCN channels and their regulatory auxiliary subunit, TRIP8b (Han et al., 2015; Saponaro et al., 2018), as well as antisense oligonucleotide-based strategies aimed at HCN1 mRNA downregulation. Of note, the latter strategy may be particularly effective in light of the hypothesis that protein misfolding may be an important component of the pathology in *HCN1*-linked DEE, together with the observation that *Hcn1* null mice do not have spontaneous seizures (Huang et al., 2009; Santoro et al., 2010).

In conclusion, the availability of strong preclinical models, such as the *Hcn1^G380D/+^* and *Hcn1^M142I/+^* mouse lines presented here, provides an ideal platform for further investigation of mechanisms underlying epileptic encephalopathies, and for testing new and paradigm-shifting therapeutic strategies.

## Acknowledgements

We acknowledge the support of animal facilities for excellent mouse care (Cologne: Esther Mahabir-Brenner (CMMC team), Maria Guschlbauer (Medical Faculty team), Branko Zevnik (CECAD in vivo Research Facility); New York: Christine Winnicker (ZMBBI). We are grateful for the expert assistance of the Transgenic Core Unit of the CECAD in vivo Research Facility (Branko Zevnik) and the Genetically Modified Mouse Model Shared Resource (GMMMSR) at Columbia University (Chyuan-Sheng Victor Lin) for the generation of transgenic mice. We thank Daniel Bauer for providing PDB files and advice on *HCN1* variant structural modeling, and Chris Reid, Nick Poolos, Carla Marini and Wayne Frankel for helpful discussion during the course of the study.

This work was supported by grants from the German Research Foundation (DFG, IS63/10-1 (FOR 2715: *Epileptogenesis of genetic epilepsies*) to D.I.; Telethon award GGP20021 to A.M.; NIH grants NS106983 and NS109366 to S.A.S.; NIH CCSG grant NCI 5P30CA013696-44 and the Columbia Precision Medicine Initiative for the generation of mouse models of human disease.

## Materials and Methods

### Animals

Mouse colonies were maintained both at the University of Cologne and at Columbia University in New York. For animals housed in Cologne, mice were kept in type II long plastic cages under standard housing conditions (21±2°C, 50% relative humidity, food (ssniff Spezialitäten GmbH, Soest, Germany) and water *ad libitum*; individually ventilated cages or ventilated cabinets (SCANBUR) and an inverted 12:12 dark:light cycle (with light turning on at 10 pm). All experiments were in accordance with European, national and institutional guidelines and approved by the State Office of North Rhine-Westphalia, Department of Nature, Environment and Consumer Protection (LANUV NRW, Germany). For animals housed in New York, mice were maintained on a 12 h light–dark cycle (with light turning on at 7 am) under standard housing conditions as above, with *ad libitum* access to food (Pico Lab rodent diet 5053 for general maintenance or Pico Lab mouse diet 5058 for breeders; Lab Diet, St. Louis, MO) and water. Weanlings were supplemented with DietGel Recovery (ClearH2O, Westbrook, ME) for two weeks post-weaning. All animal experiments were conducted in accordance with policies of the NIH Guide for the Care and Use of Laboratory Animals and the Institutional Animal Care and Use Committee (IACUC) of Columbia University.

The following commercially available mouse lines were used: B6.129P2-*Pvalb^tm1(cre)Arbr^*/J (Jackson Laboratories stock 017320; Bar Harbor, ME) and B6.129S-*Hcn1^tm2Kndl^*/J (Jackson Laboratories, stock 016566). All animals were maintained on a C57BL/6J background.

*Mouse genome editing:*

We employed CRISPR/Cas9 genome editing using the Easy Electroporation of Zygotes (EEZy) approach in order to generate *Hcn1^G380D^* (C57BL/6J-*Hcn1^em1(G380D)Cecad^*, further referred to as *Hcn1^G380D^*) and *Hcn1*^M142I^ (C57BL/6J-*Hcn1^em2(M142I)Cecad^*, further referred to as *Hcn1^M142I^*) mice as previously described (Tröder et al., 2018). gRNAs were selected using CRISPOR (Haeussler et al., 2016). *Hcn1^G380D^* mice were generated using the crRNA sequence 5’-ACTGGATCAAAGCTGTGGCA-3’ and the ssODN sequence 5’ –CCCAAGCCCCTGTCAGCATGTCTGACCTCTGGATTACCATGCTGAGCATGATTGTGGGCGCCACCTGCTACGCAATGTTTGTTGATCATGCCACAGCTTTGATCCAGTCTTTGGACTCTTCAAGGAG – 3’, containing a new Bcl1 restriction site. *Hcn1*^M142I^ mice were generated using the crRNA sequence 5’ – ATCATGCTTATAATGATGGT – 3’ and the ssODN sequence 5’ – ATCGGATGCCACGTTGAAAATAATCCACGGTGTTGTCGTCTGCTCTGTGAAGAACGTGATTCCAACTGGTATGATGACCAAATTTCCAACGATCATTATAAGCATGATTAAATCCCAATAAAACCTA – 3’, containing a new DpnII restriction site. Custom crRNAs (Alt-R), generic tracrRNAs and ssODN (<u>Ultramers)</u> were purchased from Integrated DNA Technologies (Coralville, IA, USA). Genome editing was performed in C57BL/6J mice at the *in vivo* Research Facility of the CECAD Research Center, University of Cologne, Germany. Identical procedures were used to generate *Hcn1^G380D^* and *Hcn1^M142I^* mice in the C57BL/6J background at the Genetically Modified Mouse Models shared resource of Columbia University, New York. All founder lines (two for *Hcn1^G380D^* and two for *Hcn1^M142I^*) were assessed for basic phenotypic features (growth, brain size, presence of spontaneous seizures, pathohistological markers, HCN1 protein expression, drug response) with no differences noted.

### Mouse genotyping

DNA was isolated from ear biopsies in 100 µl lysis buffer (100 mM NaCl, 50 mM Tris/HCl pH 8.0, 1 mM EDTA, 0.2% Nonidet P-40, 0.2% Tween 20, 0.1 mg/ml Proteinase K) over night at 54°C under constant shaking, followed by 40 min at 84°C to inactivate the proteinase K. Amplification was performed in PCR reaction mix using a touchdown PCR protocol with the following primers: *Hcn1^G380D^* wildtype (WT) forward 5’ – ACGGTGATGACACTTGTTCAGT – 3’, reverse 5’ – TGGATCAAAGCTGTGGCATGGC – 3’ (468 bp); *Hcn1^G380D^* mutant forward 5’ – ACCTGCTACGCAATGTTTGTTGAT – 3’, reverse 5’ – GGCACTACACGCTAGGAA 3’ (354 bp); *Hcn1^M142I^* forward 5’ – CAACATTTGTTTGTTCTCCTCACC – 3’, reverse 5’ – ATGATCGAATGCCACGTTGA – 3’ (248 bp). For *Hcn1^M142I^*, a restriction digest followed to identify heterozygous animals containing the DpnII restriction site. DNA purification was performed with GeneJET PCR Purification kit (Thermo Scientific, Germany) according to manufacturer’s instructions. Enzyme restriction with DpnII was performed in 30 µl reaction mix (1 µl DpnII, 3 µl DpnII buffer, 10 µl purified DNA and 16 µl H_2_O; New England Biolabs, Ipswich, MA) and incubation at 37°C for 2h. Gel electrophoresis was performed with 1.8% agarose gels (VWR Life science, Sigma-Aldrich) in 1X TAE buffer (40 mM Tris, 10 mM acetic acid, 1 mM EDTA pH 8.0) using a 200 bp DNA ladder (PANladder, PAN-Biotech™). Genotyping for each animal was performed in two technical replicates at the beginning and end of the experiments, respectively.

### Electrocorticogram (ECoG) recordings

Surgery: Adult animals of both genotypes and sexes were implanted with radio transmitters (PhysioTel^®^ ETA-F10, Data Sciences International) for long-term electrocorticogram (ECoG) and video monitoring. For analgesia, mice received 0.025 mg/kg buprenorphine (i.p., TEMGESIC®, Indivior, UK Limited) prior to surgery and 5.0 mg/kg Carprofen (s.c., Norbrook® Laboratories Limited, Ireland) during surgery. Mice were anesthetized with 0.5 – 4% isoflurane in 100% oxygen, and kept at 0.8 – 1.5% isoflurane throughout the surgery. Body temperature was maintained at 36.5°C using a homeothermic heating pad (Stoelting, Germany). Mice were placed into a stereotaxic device (Kopf instruments, CA, USA), a midline skin incision was made above the skull and the periosteum was denatured by a short treatment with 10% H_2_O_2_, followed by rinsing with 0.9% NaCl solution, drying of the skull and application of dental cement (OptiBond™, Kerr Dental, Germany) to harden the skull surface. The transmitter body was implanted subcutaneously in a pouch made in the loose skin of the back. The two lead wires were tunneled subcutaneously through the incision on the skull. Using a dentral drill, a small hole was drilled above the hippocampus (2 mm posterior from Bregma, 2 mm from the midline, always on the right side) for the recording wire, and another hole was made above the cerebellum for the reference wire. The wires were placed into the holes so as to touch the dura and fixed with dental cement. The skin was subsequently closed with tissue glue (GLUture^®^, World Precision Instruments, WPI, USA) and animals were allowed to recover for 5 days before data acquisition.

Data acquisition and analysis: Recordings were performed by placing the animal’s home cage onto a receiver board that detected the ECoG and movement activity signals, and together with the synchronized video recordings data were digitally stored using Ponemah (DSI™, MN, USA) and the Media Recorder (Noldus, Wageningen, The Netherlands). Seizure and spike detection was performed with the software NeuroScore (DSI™, MN, USA) using an integrated, automated spike detection tool (absolute threshold: threshold value 200 µV, maximum value 2000 µV; spike duration: 0.1 ms – 250 ms) and manually verified by the experimenter. Graphical visualization of the seizures was performed with a custom script in MATLAB® (MathWorks, MA, USA).

### Behavioral analysis

Behavioral grading of seizures: For evaluation of seizures, animals were monitored electrographically and behaviorally and seizures graded using a modified Racine scale (Van Erum et al., 2019), with 0 = normal behavior; 1 = mouth and facial movements; 2 = head nodding; 3 = forelimb clonus; 4 = rearing with forelimb clonus, falling; 5 = wild running, jumping; 6 = laying on the side with fore- and hindlimb clonus. When multiple classes of severity occurred during one electrographically defined seizure, the most severe behavior represented the grade of that seizure.

Open field: The open field was performed in a box (50 x 50 x 40 cm) illuminated with 100 lux. Mice were placed in one corner of the arena and could freely move for 15 min. Tracks were recorded and analyzed with the software EthoVision (Noldus, Wageningen, The Netherlands). The following parameters were obtained: distance moved, running velocity, number of rotations, time in border (an imaginary 5 cm wide border around the arena), and time in center (an imaginary inner square of 20 x 20 cm). All experiments were performed during the dark cycle when animals are naturally active.

Gait analysis: For the automated gait analysis system Catwalk XT (Noldus, Wageningen, The Netherlands) mice had to run on an enclosed walkway on a glass plate. Run duration variation was set to 0.5-20 sec, with a maximum speed variation of 60%, and a minimum of ten consecutive steps. A minimum of eight runs per mouse was collected. Runs were classified with the Catwalk XT software (Noldus, Wageningen, The Netherlands). Footprints were detected automatically by the software and manually corrected by visual inspection. The following parameters were used for analysis: running speed, stand (duration in seconds of contact of a paw with the glass plate), stride length (distance between successive placements of the same paw in centimeter), step cycle (the time in seconds between two consecutive paw placements), base of support (BOS, the average width between the hind paws), step sequence (contains information on the order in which four paws are placed), and regularity index (expresses the number of normal step sequence patterns relative to the total number of paw placements in percent).

### Drug testing

Lamotrigine (23 mg/kg lamotrigine isethionate; Tocris, Minneapolis, MN) and valproate (250 mg/kg sodium valproate; Tocris) were dissolved in saline (0.9% NaCl). Phenytoin (30 mg/kg Phenhydan; Desitin, Hamburg, Germany) was diluted in 21% cyclodextran/ 1X PBS (pH 8.0) or administered as fosphenytoin (45 mg/kg fosphenytoin sodium, corresponding to 30 mg/kg phenytoin equivalents or PE; Millipore-Sigma, St. Louis, MO) dissolved in saline. Saline and 21% cyclodextran/ 1X PBS, respectively, were used as vehicle. All drugs were applied by intraperitoneal injection (i.p.). For behavioral seizure assessment, animals were observed for a total of 4 h after injection. For video- ECoG seizure assessment, animals were recorded for 24 h after injection. The highest severity seizure displayed during the observation period was scored as the seizure grade for that animal’s response to drug administration.

### Immunohistochemistry

Animals were perfused with 1X phosphate buffered saline (PBS) followed by 4% paraformaldehyde in 1X PBS, and brains post-fixed overnight at 4 °C. After several washes in 1X PBS, 40 µm coronal slices were cut using a vibratome, and free floating sections permeabilized in PBS + 0.1% Triton, followed by incubation in blocking solution (1X PBS + 5% normal donkey serum) for 1 h at room temperature. Primary antibody incubation was carried out in blocking solution overnight at 4°C. Antibodies used were: mouse monoclonal anti-HCN1 (clone N70-28, NeuroMab 75-110, dilution 1:300; Davis, CA); rabbit polyclonal anti-NPY (ImmunoStar 22940, dilution 1:1000; Hudson, WI); mouse monoclonal anti-GFAP (clone GA5, Invitrogen 14-9892-82, dilution 1:250); mouse monoclonal anti-ATP1a3 (clone G10, Invitrogen MA3-915, dilution 1:400); rat monoclonal anti-HA tag (clone BMG-3F10, Roche 12013819001, dilution 1:100); mouse monoclonal anti-K_v_1.2 (clone K14/16, NeuroMab 75-008, dilution 1:300). Secondary antibody incubation was performed in blocking solution for 2 h at room temperature. All secondary antibodies were used at 1:500 dilutions: goat anti-mouse IgG1 cross-adsorbed (Alexa Fluor 488, Life Technologies A21121; Eugene, OR), donkey anti-rabbit cross-adsorbed (Alexa Fluor 488, Life Technologies A21206), goat anti-rabbit Superclonal (Alexa Fluor 647, Life Technologies A27040), goat anti-rat IgG (H+L) cross-adsorbed (Alexa Fluor 488, Life Technologies, A1106) and goat anti-mouse IgG2b cross-adsorbed (Alexa Fluor 488, Life Technologies, A21141). For perineuronal net staining, WFA biotin conjugate was used (Sigma L1516, dilution 1:1000), followed by incubation with Streptavidin (Alexa Fluor™ 594 conjugate, Life Technologies S32356, dilution 1:500). Nissl stain was performed by adding NeuroTrace reagent (NT 640/660, Thermofisher N21483, dilution 1:500) during the secondary antibody incubation, followed by extensive washing in 1X PBS. Images were acquired on a Zeiss LSM 700 laser scanning confocal microscope with Zen 2012 SP5 FP3 black edition software, using either a Zeiss Fluar 5x/0.25 objective (0.5 zoom, pixel size: 2.5 x 2.5 µm^2^) or a Zeiss Plan-Apochromat 20X/0.8 objective (1.0 zoom, pixel size: 0.3126 x 0.3126 µm^2^). Image analysis and fluorescent signal quantification were performed using ImageJ 1.49v software (National Institutes of Health, USA).

### Stereotaxic virus injection

Mice were anaesthetized using isoflurane (Covetrus, Portland, ME) and provided analgesics (Carprofen, Zoetis, Troy Hills, NJ). A craniotomy was performed above the target region and a glass pipette was stereotaxically lowered to the desired depth. Injections were performed using a nano-inject II apparatus (Drummond Scientific), with 25 nl of solution delivered every 15 s until a total amount of 200 nl was reached. The pipette was retracted after 5 min. Viruses were injected bilaterally, at a titer of 2x10^12^ cfu/µl, with injection coordinates AP –2.0, ML ±1.5, DV –1.4 (in millimetres with Bregma as reference).

For virus construction, Addgene plasmid #44362 (pAAV-hSyn-DIO-hM4D(Gi)- mCherry) was cut with restriction enzymes NheI and AscI, and the hM4D(Gi)-mCherry cDNA was replaced with mouse HCN1 cDNA including an N-terminal HA tag (YPYDVPDYA) immediately following the starting methionine. Site-directed mutagenesis was performed by Applied Biological Materials Inc. (Richmond, BC, Canada) and viral DNA packaged into adeno associated virus (AAV) serotype 8 by the Duke University Viral Vector Core (Durham, NC).

### Slice electrophysiology

Slice preparation: Mice were anesthetised by inhalation of isoflurane (5%) for 7 minutes, subjected to cardiac perfusion of ice-cold carbogenated artificial cerebrospinal fluid, modified for dissections (d-ACSF; 195 mM sucrose, 10 mM glucose, 10 mM NaCl, 7 mM MgCl_2_, 0.5 mM CaCl_2_, 25 mM NaHCO_3_, 2.5 mM KCl, 1.25 mM NaH_2_PO_4_, 2 mM Na-pyruvate, pH 7.2) for 30 s before decapitation according to the procedures approved by the IACUC of Columbia University. The skull was opened, and the brain removed and immediately transferred into ice-cold carbogenated d-ACSF. The hippocampus was dissected in both hemispheres. Each hippocampus was placed in the groove of an agar block and 400 µm thick hippocampal slices were cut using a vibrating tissue slicer (VT 1200, Leica, Germany) and transferred to a chamber containing a carbogenated mixture of 50% d-ACSF and 50% ACSF (22.5 mM glucose, 125 mM NaCl, 1 mM MgCl_2_, 2 mM CaCl_2_, 25 mM NaHCO_3_, 2.5 mM KCl, 1.25 mM NaH_2_PO_4_, 3 mM Na-pyruvate, 1mM ascorbic acid, pH 7.2) at 35°C, where they were incubated for 40 - 60 min. Thereafter, slices were held at room temperature (21°C) until transfer into the recording chamber.

Electrophysiology: Slices were transferred from the incubation chamber into the recording chamber of an Olympus BX51WI microscope (Olympus, Japan), where they were held in place by a 1mm grid of nylon strings on a platinum frame. Slices were continually perfused with ACSF at 34 ± 1°C, maintained by a thermostat-controlled flow- through heater (Warner Instruments, CT, USA). Healthy somas of CA1 pyramidal neurons were identified visually under 40X (20 x 2) magnification and patched under visual guidance using borosilicate glass pipettes (I.D. 0.75 mm, O.D. 1.5 mm, Sutter Instruments, UK) with a tip resistance of 4 – 5.5 MΩ, connected to a Multiclamp 700B amplifier (Molecular Devices, CA) and filled with intracellular solution, containing (in mM): 135 K-gluconate, 5 KCl, 0.1 EGTA, 10 HEPES, 2 NaCl, 5 MgATP, 0.4 Na_2_GTP, 10 Na_2_-Phosphocreatin, adjusted to a pH of 7.2 with KOH. Recordings were only accepted if the series resistance after establishing a whole-cell configuration did not exceed 25 MΩ and did not change by more than 20% of the initial value during the course of the experiment.

Pharmacology: Stock solution of 10 mM ZD7288 (Tocris, UK) was stored at -20 °C and diluted in ACSF to a concentration of 10 µM before bath application to the slice. In some experiments, blockers of GABA_A_ and GABA_B_ receptors, 2 μM SR95531 and 2 μM CGP55845, respectively, were added to the bath solution. As application of the latter did not alter any of the reported measures and since synaptic transmission was not assessed in this case, data were pooled between experiments performed with and without blockers of inhibition.

Data acquisition and analysis: Electrophysiological recordings were digitized, using a Digidata 1322A A/D interface (Molecular Devices, CA), at a sampling rate of 20 kHz (low pass filtered at 10 kHz) and recorded pClamp 10 software (Molecular Devices, CA, USA). The amplifier setting of the Multiclamp 700B were controlled through Multiclamp Commander (Molecular Devices, CA, USA).

After 50 ms baseline recording, 1-s current steps of –350 to +350 pA were applied to the patched cells in increments of 25 pA, after which an additional 1s of post-step membrane potential was recorded. The trigger time between these episodes was 3 s. Voltage deflections in response to current steps of –50 to +50 pA were used to calculate the input resistance. Initial resting membrane potential (RMP) was obtained immediately upon breaking into the cell. Voltage sag in response to negative current steps was calculated by dividing the steady state voltage deflection during the late phase of the –100 pA current step by the peak of the voltage deflection during the same step.

Data was analyzed using Axograph X software (Axograph Scientific, Australia), MATLAB (Mathworks, MA, USA) as well as Microsoft Excel (Microsoft Corp., WA, USA) or Prism 8 (Graphpad, CA, USA) and visualized in Acrobat Illustrator (Adobe, CA, USA).

### HEK293 cell electrophysiology

Constructs: The cDNAs encoding full-length human HCN1 channel and mouse TRIP8b (splicing variant 1a-4) were cloned into the pcDNA 3.1 (Clontech Laboratories) mammalian expression vector. The cDNA encoding the full-length mouse HCN2 was cloned in pCI (Promega) mammalian expression vector. The cDNA encoding the full- length human Na_v_1.5 channel was cloned in pIRES-EGFP mammalian expression vector. Cell culture and transfection: HEK293T cells were cultured in Dulbecco’s modified Eagle’s medium (Euroclone) supplemented with 10% fetal bovine serum (Euroclone), 1% Pen Strep (100 U/ml of penicillin and 100 µg/ml of streptomycin) and grown at 37°C with 5% CO2. When ∼70% confluent, HEK293T cells were transiently transfected with cDNA using Turbofect transfection reagent (Thermo Fisher, Germany) according to the manufacturers recommended protocol. For each 35 mm Petri dish, 1 µg of the plasmid DNA and 0.3 µg of EGFP-containing vector (pmaxGFP, AmaxaBiosystems) were used. For HCN1 and TRIP8b coexpression, 1 µg of each cDNA was used in combination with 0.3 µg of EGFP-containing vector.

Chemicals: Lamotrigine isethionate (Tocris) and ZD7288 (Tocris) stock solutions were prepared by dissolving the powders in milliQ water to obtain a final stock concentration of 100 mM and 10 mM, respectively. Single-use aliquots were made and stored at −20°C until the day of the experiment.

Electrophysiology and data analysis: 30 – 72 h after transfection, the cells were dispersed by trypsin treatment. Green fluorescent cells were selected for patch-clamp experiments at room temperature (about 25°C). Currents were recorded in whole-cell configuration either with a ePatch amplifier (Elements, Cesena, Italy) or with a Dagan3900A amplifier (Dagan Corporation, MN, USA); data acquired with the Dagan3900A amplifier were digitized with an Axon Digidata 1550B (Molecular Devices, CA, USA) converter. All data were analysed off-line with Axon pClamp 10.7. Patch pipettes were fabricated from 1.5 mm O.D. and 0.86 I.D. borosilicate glass capillaries (Sutter, Novato, CA, USA) with a P-97 Flaming/Brown Micropipette Puller (Sutter, Novato, CA, USA) and had resistances of 3 - 6 MΩ. For the recordings of HCN channels, the pipettes were filled with a solution containing: 10 mM NaCl, 130 mM KCl, 1 mM egtazic acid (EGTA), 0.5 mM MgCl_2_, 2 mM ATP (Magnesium salt) and 5 mM HEPES–KOH buffer (pH 7.2), while the extracellular bath solution contained 110 mM NaCl, 30 mM KCl, 1.8 mM CaCl_2_, 0.5 mM MgCl_2_ and 5 mM HEPES–KOH buffer (pH 7.4). For the recordings of Na_v_1.5 channel, the pipettes were filled with a solution containing 5 mM NaCl, 140 mM CsCl, 4mM ATP (Magnesium salt), 2 mM MgCl_2_, 5 mM EGTA, 10 mM HEPES-CsOH buffer (pH 7.4) while the extracellular bath solution contained: 135 mM NaCl, 4mM KCl, 1 mM CaCl_2_, 2 mM MgCl_2_, 20 mM D-glucose, 10 mM HEPES-NaOH buffer (pH 7.4). To assess HCN channel activation curves, different voltage-clamp protocols were applied depending on the HCN subtype: for HCN1 and HCN1 coexpressed with TRIP8b, holding potential was –20 mV (1 s), with steps from –30 mV to –120 mV (–10 mV increments, 3.5 s) and tail currents recorded at –40 mV (3.5 s); for HCN2, holding potential was –20 mV (1 s), with steps from -40 mV to -130 mV (-15 mV increments, 5 s) and tail currents recorded at –40 mV (5 s). Patch-clamp currents were acquired with a sampling rate of 5 kHz and lowpass filtered at 2.5 kHz. Na_v_1.5 recordings were obtained from holding potentials of -80 mV or -130 mV, as indicated. Currents were elicited by stepping to -30 mV for 50 ms followed by a step to the next holding potential (–60 or –130 mV for 5 s) in a cycling manner. To measure the inactivation curves a voltage step protocol was used starting from a holding potential of –100 mV for 150 ms followed by a series of inactivating pulses (from –30 to –130 mV, for 500 ms); the fraction of channels which remain available after each inactivating pulse were assessed by the peak currents during the following short test pulse at 0 mV for 50 ms. LTG was applied to the (extracellular) bath solution of the experiments at 100 µM concentration. For HCN channels, the effect was assessed after 20 min of application of LTG in the bath solution. The percentage of ZD7288 induced block (30 µM concentration, bath applied) was assessed by measuring the current amplitude at the end of the hyperpolarizing pulse at -110 mV, which includes both the instantaneous and the time dependent component of the current. After reaching the full effect of ZD7288, a 5 mM CsCl-containing solution was added to the bath to verify current response to a second known HCN channel blocker. Mean activation curves were obtained by fitting maximal tail current amplitude, plotted against the voltage step applied, with the Boltzmann equation:

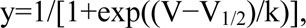

where V is voltage, y the fractional activation, V_1/2_ the half-activation voltage, and k the inverse-slope factor = –RT/zF (all in mV), using Originpro software (Originlab, Northampton, MA, USA). Mean V_1/2_ values were obtained by fitting individual curves from each cell to the Boltzmann equation and then averaging all the obtained values. All measurements were performed at room temperature.

### Data analysis

Sample size estimation was based on prior studies and experience (Festing, 2018). For behavioral experiments in Figure 2, sample size was calculated using a power analysis assuming an effect size of 0.5, a power of 0.8 and an α error of 0.05, based on previous experience (G*Power 3.1.9.2, University Düsseldorf, http://www.gpower.hhu.de). No data were excluded after analysis. Statistical analysis was performed using GraphPad Prism (Version 9.0.1, Graphpad, CA, USA). Parameters were assessed for normality using the D’Agostino & Pearson test. For normally distributed data, means were compared using a two-tailed Students *t* tests (assuming equal variances between genotypes) and paired data was analysed with a paired *t* test. For repeated measurements, a two-way repeated measurement (RM) ANOVA was performed, and, when appropriate, a *post hoc* Šídák’s multiple comparisons test followed. For non-parametric data, medians were compared using the Mann Whitney *U* test, paired data were analysed with a Wilcoxon matched-pairs signed rank test, and in case of two grouping factors a Kruskal-Wallis test with *post hoc* Dunn’s multiple comparisons, when appropriate, was performed. All tests were two-tailed, and statistical significance accepted at *P* < 0.05. All parameters were assessed for a sex difference with a mixed-effects analysis, having *genotype* and *sex* as grouping factors. Since we did not find sex-specific differences (except for parameters shown in Figures 1A and 1C), data from female and male subjects were pooled. Unless otherwise stated, data represent mean ± SEM.

---

Video 1. Video-ECoG recording of a spontaneous seizure in *Hcn1^G380D/+^* heterozygous female. The first portion of the video illustrates a period of grade 3-4 behavioral seizures, accompanied by high amplitude oscillatory activity on the ECoG trace. Immediately following the high amplitude ECoG activity, note the onset of post-ictal depression (start at 1:33:36 time mark on EcoG trace, approximately) as indicated by the flat ondulatory trace on ECoG and accompanying behavioral immobility.

Video 2. Video-ECoG recording of a spontaneous seizure in *Hcn1^M142I/+^* heterozygous male. The first portion of the video illustrates grade 2-5 behavioral seizures, accompanied by high amplitude oscillatory activity on the ECoG trace, similar to *Hcn1^G380D/+^* animals. This is followed by a period of irregular, low amplitude cortical activity on the ECoG trace as seizure activity moves into lower brain areas, accompanied by grade 6 behavioral seizures (start at 12:04:31 time mark on EcoG trace, approximately). Post-ictal depression can be noted in the last few seconds of the clip.

**Figure 1 — figure supplement 1:**
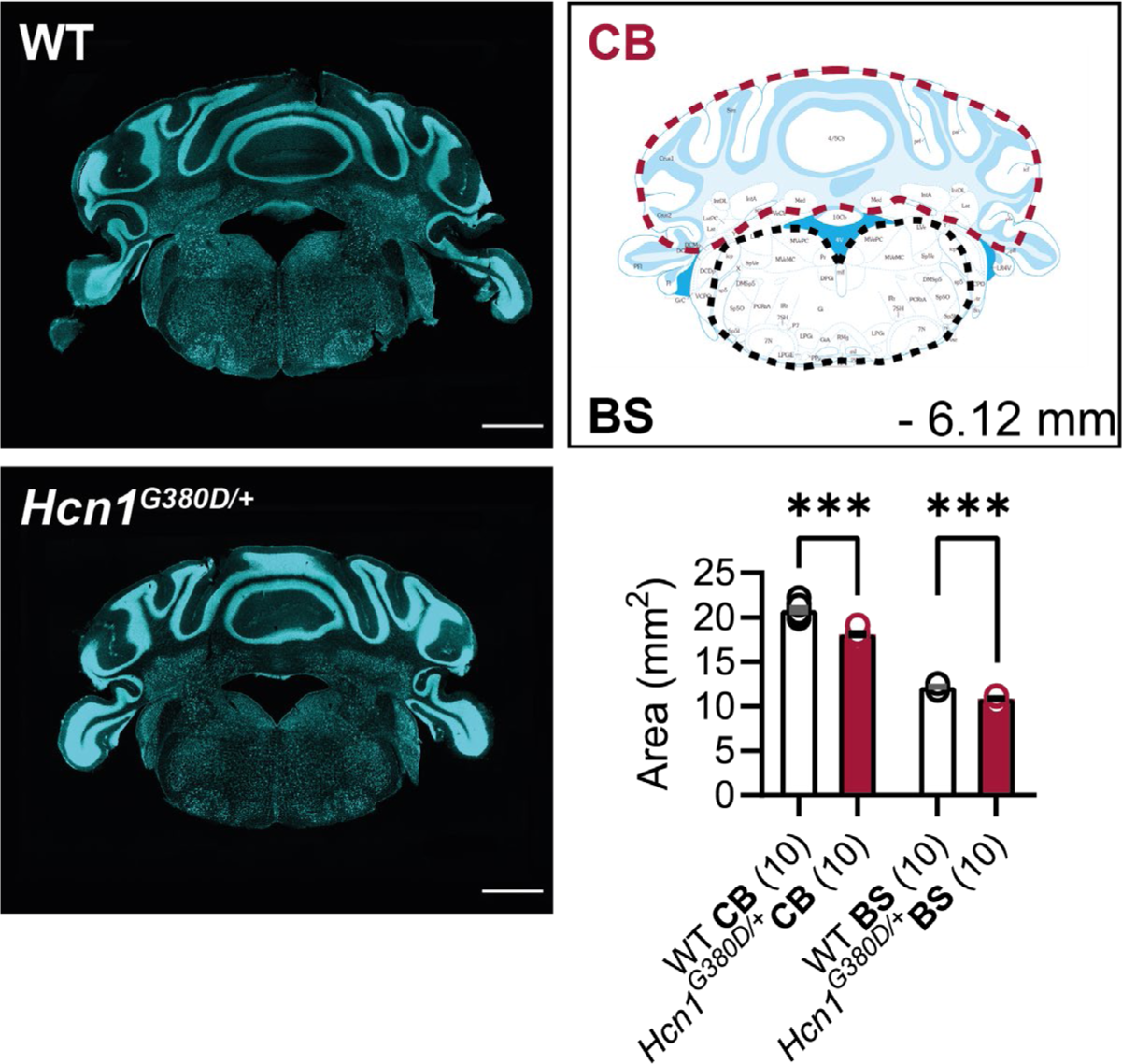
Cerebellum and brainstem area in *Hcn1^G380D/+^* mice. Fluorescent Nissl stain of hindbrain sections from WT and *Hcn1^G380D/+^* mice are shown on the left (Bregma –6.12 mm, scale bar = 1200 μm). Brain area measurements comparing cerebellum (CB) and brainstem (BS) shown on the right (CB: WT 20.84 ± 0.23 mm^2^, *Hcn1^G380D/+^* 18.11 ± 0.19 mm^2^; BS: WT 12.18 ± 0.06 mm^2^, *Hcn1^G380D/+^* 10.87 ± 0.11 mm^2^). Cerebellum area was more affected than brainstem area (2-way RM ANOVA, *brain region* x *genotype* interaction F(1, 9) = 49.56, *** *P* < 0.0001), with mutant cerebellum 13.1% smaller and mutant brainstem 10.8% smaller, compared to WT. Measured areas are indicated by dotted lines (for cerebellum measurements, flocculus and paraflocculus were excluded on both sides; for brainstem, dorsal and ventral cochlear nuclei were excluded).

**Figure 2 — figure supplement 1:**
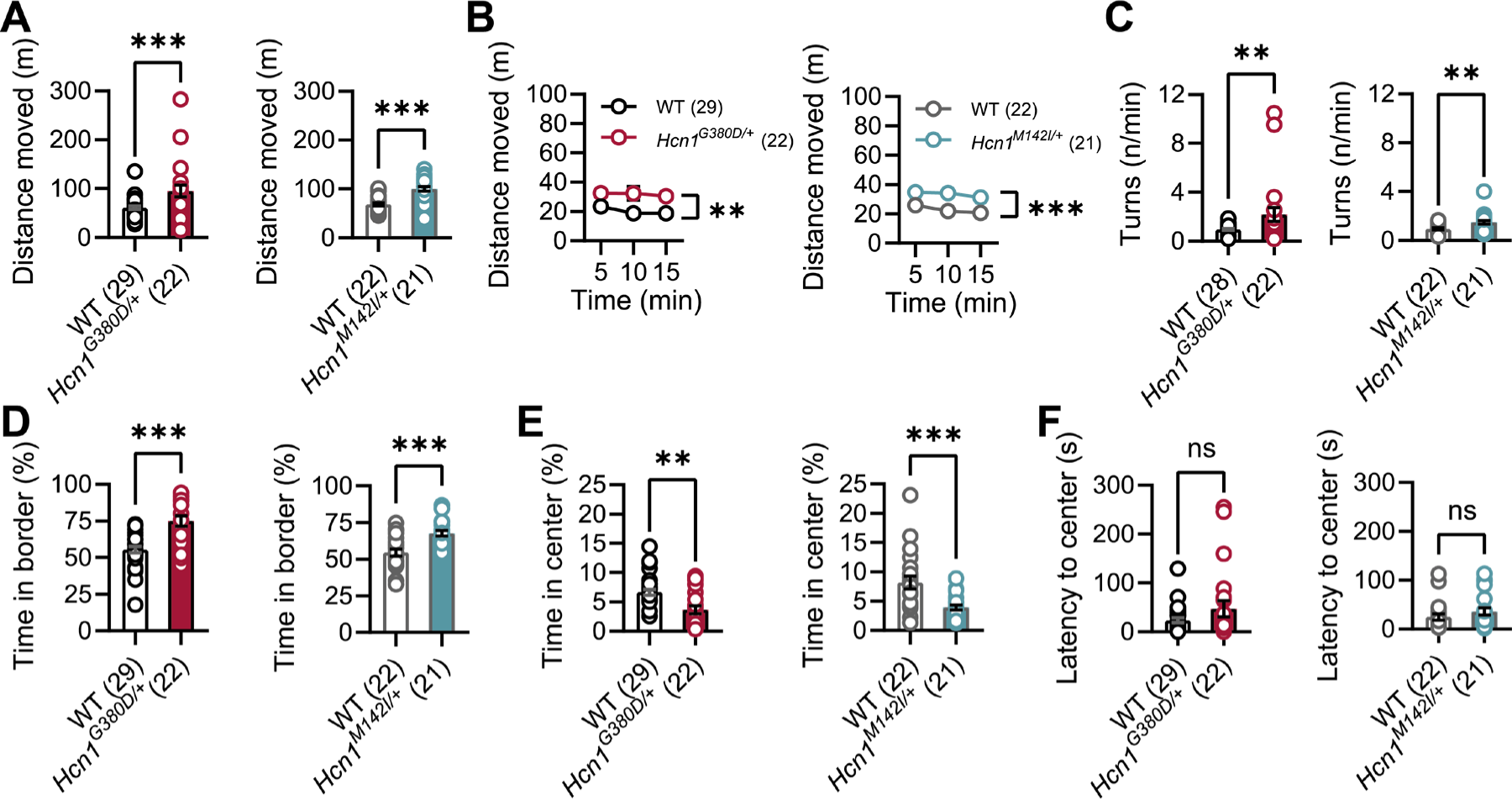
Open field behavior of *Hcn1^G380D/+^* and *Hcn1^M142I/+^* mice. Comparison of open field behavior between *Hcn1^G380D/+^* or *Hcn1^M142I/+^* mice and their respective WT littermates. A) Mean distance moved and B) mean distance moved over time (in m) during 15 min in the open field (** *P* < 0.01, *** *P* < 0.001, effect of *genotype*, 2-way RM ANOVA having *genotype* and *time* as grouping factors). C) Mean number of body rotations (turns/min). D) Mean time spent in border and E) center zones (% total trial duration). F) latency to enter the center zone (in s). (ns = not significant, ** *P* < 0.01, *** *P* < 0.001) Data represent mean ± S.E.M. (See Figure 2 – source data 1 for numerical values and statistical test used for each comparison.)

**Figure 2 — figure supplement 2:**
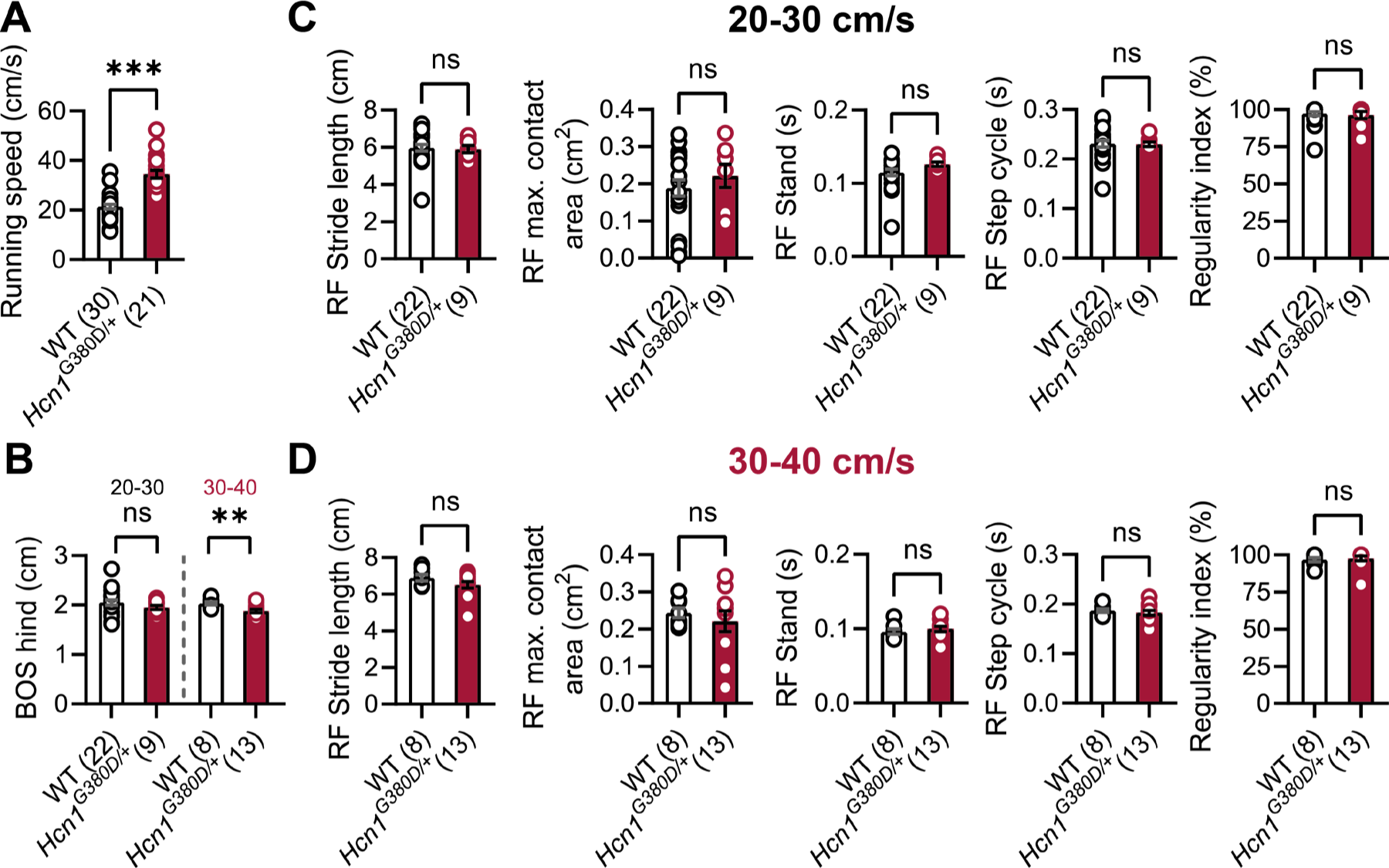
Gait analysis in *Hcn1^G380D/+^* mice. A) Mean running speed of all runs was increased in *Hcn1^G380D/+^* mice compared to WT littermates. B) BOS of the hind paws was decreased at the higher speed range in *Hcn1^G380D/+^* mice. Gait analysis parameters of the right front (RF) paw for speed ranges of C) 20-30 cm/s and D) 30-40 cm/s, including, from left to right: stride length, maximum (max.) contact area, stand, step cycle, and regularity index, were not changed in G380D knock-in mice. The same results were obtained for the other three paws (data not shown). (ns = not significant, ** *P* < 0.01, *** *P* < 0.001) Data represent mean ± S.E.M. (See Figure 2 – source data 2 for numerical values and statistical test used for each comparison).

**Figure 2 — figure supplement 3:**
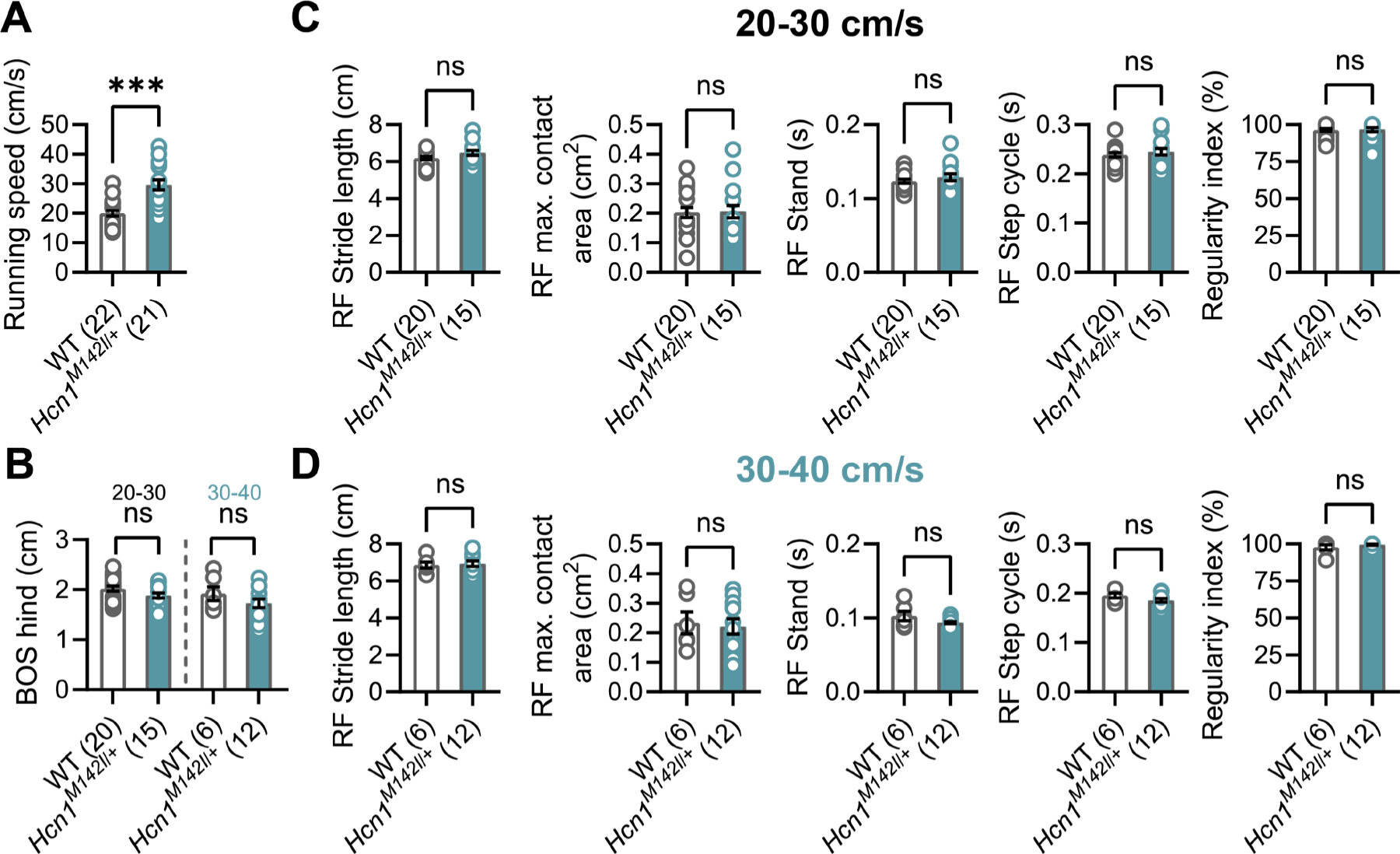
Gait analysis in *Hcn1^M142I/+^* mice. A) Mean running speed of all runs was increased in *Hcn1^M142I/+^* mice compared to WT littermates. B) BOS of the hind paws was unchanged. Gait analysis parameters of the right front (RF) paw for speed ranges of C) 20-30 cm/s and D) 30-40 cm/s, including, from left to right: stride length, maximum (max.) contact area, stand, step cycle, and regularity index. Despite significantly increased running speeds in *Hcn1^M142I/+^* mice, other gait parameters were not changed. The same results were obtained for the other three paws (data not shown). (ns = not significant, *** *P* < 0.001) Data represent mean ± S.E.M. (See Figure 2 source data 2 for numerical values and statistical test used for each comparison).

**Figure 3 — figure supplement 1:**
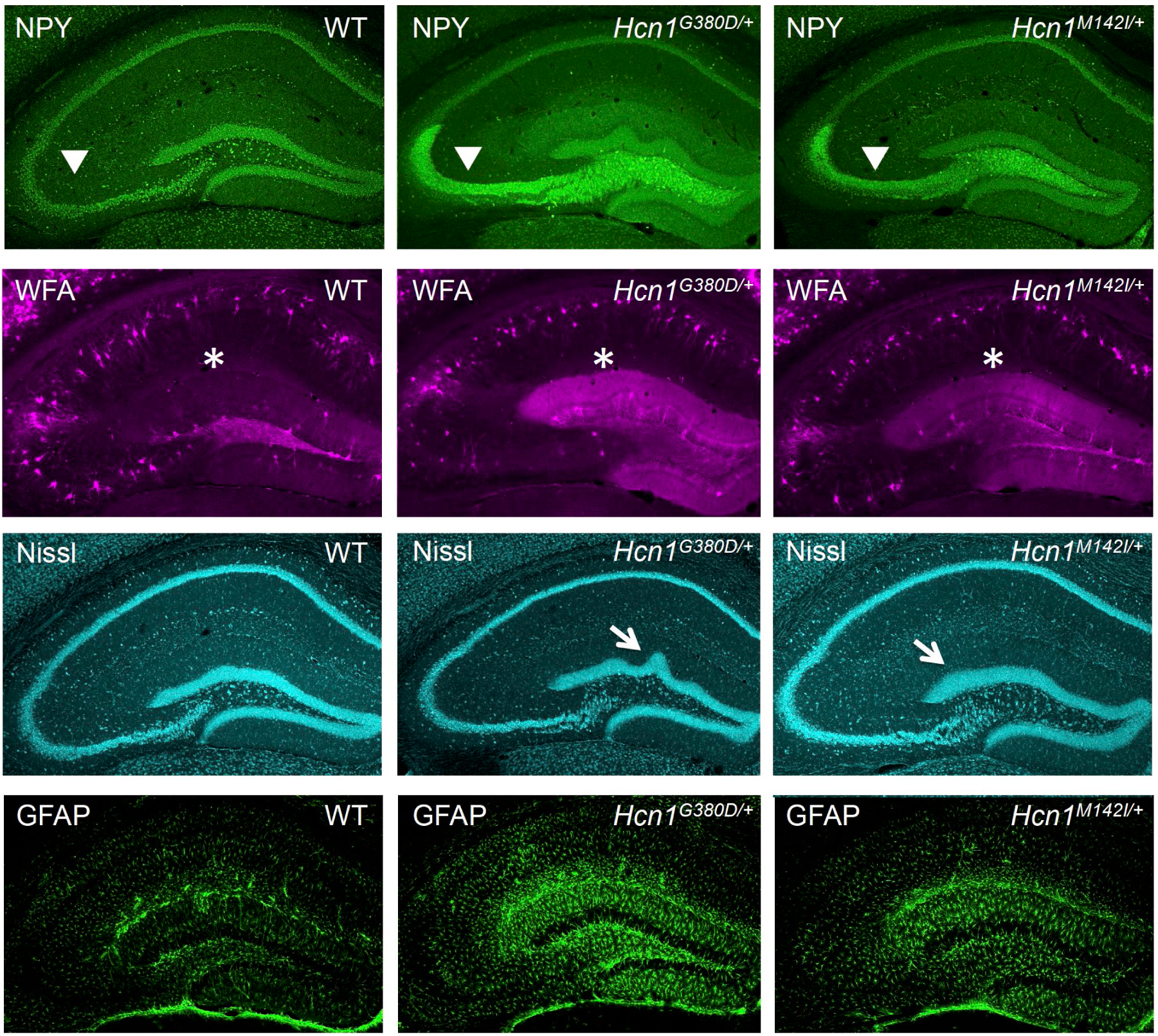
Histological markers show hippocampal changes associated with epilepsy in *Hcn1^G380D/+^* and *Hcn1^M142I/+^* mice. Immunofluorescent staining of mid-coronal sections from adult brains in WT, *Hcn1^G380D/+^* and *Hcn1^M142I/+^* mice showing hippocampal region labeled for neuropeptide Y (NPY, first row); Wisteria floribunda agglutinin (WFA, second row); Nissl bodies (Nissl, third row); and glial fibrillary protein (GFAP, bottom row). Arrowheads highlight the position of the granule cell mossy fiber tract (top row), asterisks the hippocampal fissure, located above the dentate gyrus molecular layer (second row). Note ectopic expression of NPY in mossy fiber tract, increased WFA labeling in the molecular layer of dentate gyrus, granule cell layer dispersion and upper blade gyrification (arrows in third row), and increased gliosis in mutant animals from both lines.

**Figure 4 — figure supplement 1:**
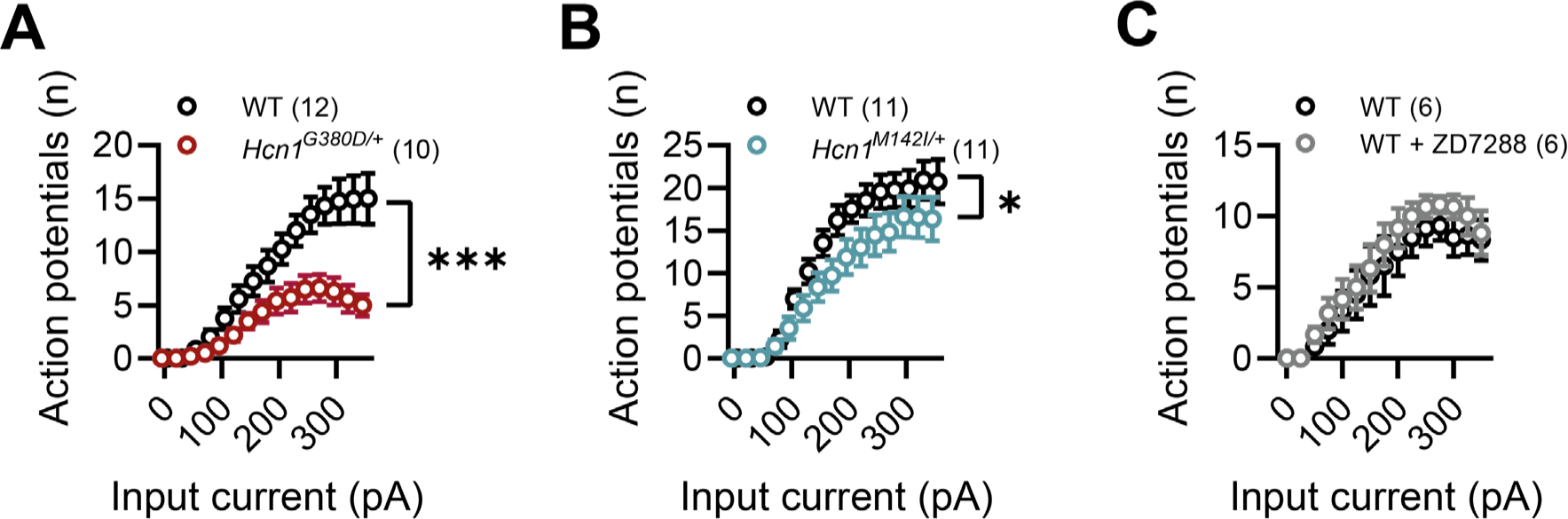
Impaired action potential firing in *Hcn1^G380D/+^* and ***Hcn1^M142I/+^* neurons**. A) Input-output curves (number of action potentials *vs* injected current) for CA1 pyramidal neurons from WT and *Hcn1^G380D/+^* littermates. B) Same comparison for CA1 pyramidal neurons from WT and *Hcn1^M142I/+^* littermates. C) Same comparison for CA1 pyramidal neurons from WT control mice, before and after bath application of 10 μM ZD7288. Note strong suppression of excitability in mutant lines not seen with acute blockade of HCN channels with ZD7288. (* *P* < 0.05, *** *P* < 0.001, effect of *genotype* x *input current*, after 2-way RM ANOVA having *genotype* and *input current* as grouping factors; number of cells is indicated in the graphs; number of animals used: G380D WT n = 4, *Hcn1^G380D/+^* n = 5; M142I WT n = 3, *Hcn1^M142I/+^* n = 3; WT ± ZD7288, n = 2). Data represent mean ± S.E.M.

**Figure 5 — figure supplement 1:**
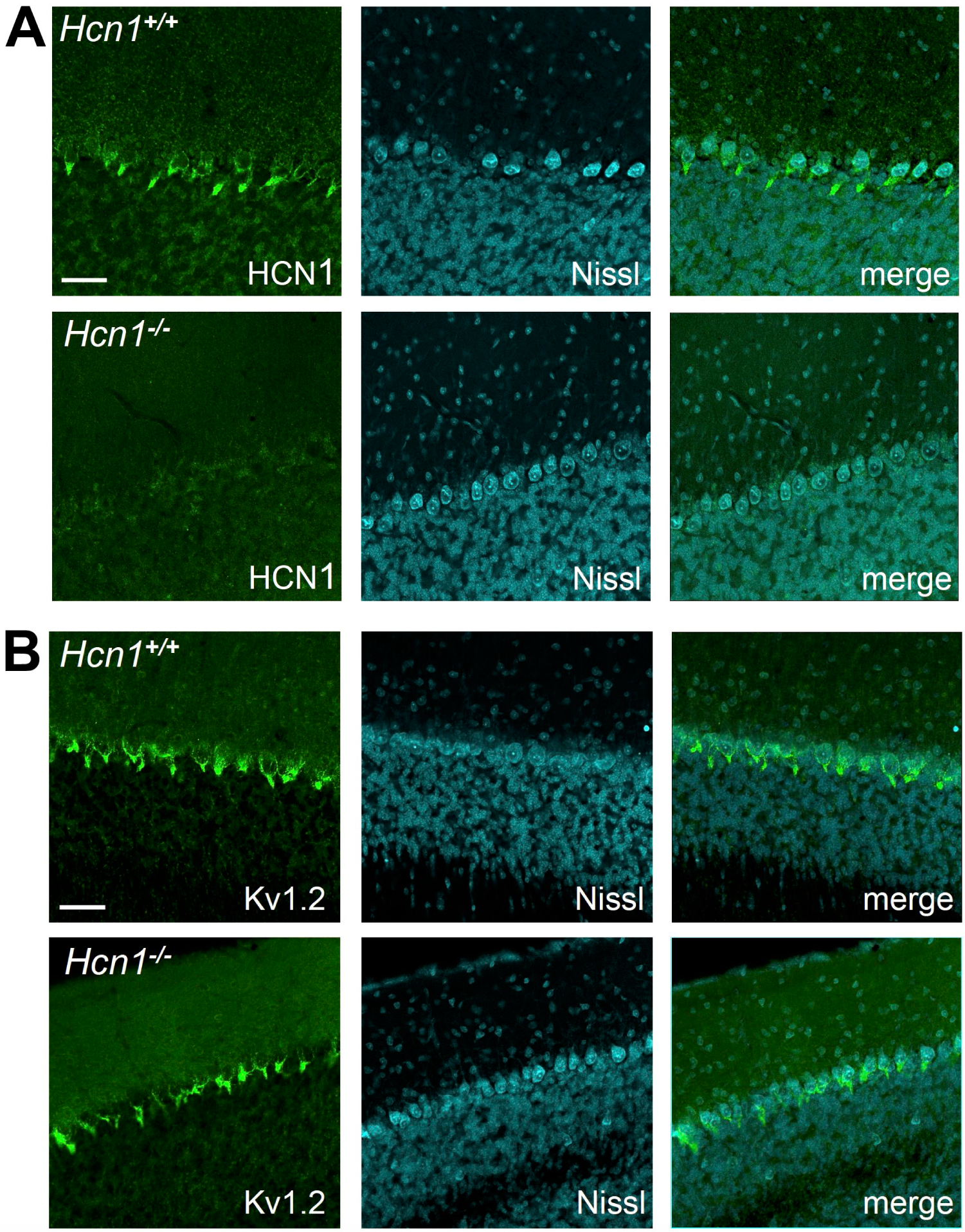
Kv1.2 protein expression is unaltered in cerebellar pinceau of global HCN1 knockout animals. Immunofluorescent staining of coronal hindbrain sections from adult brains in *Hcn1^+/+^* and *Hcn1^-/-^* mice, showing the cerebellar cortex. A) Left, HCN1 protein labeling; center, Nissl counterstain; right, merged images. B) Left, Kv1.2 protein labeling; center, Nissl counterstain; right, merged images. Nissl counterstain highlights the location of Purkinje cell bodies. The *pinceau* is visible surrounding the Purkinje cell axon initial segment. Scale bar = 50 μm in all panels.

**Figure 9 — figure supplement 1:**
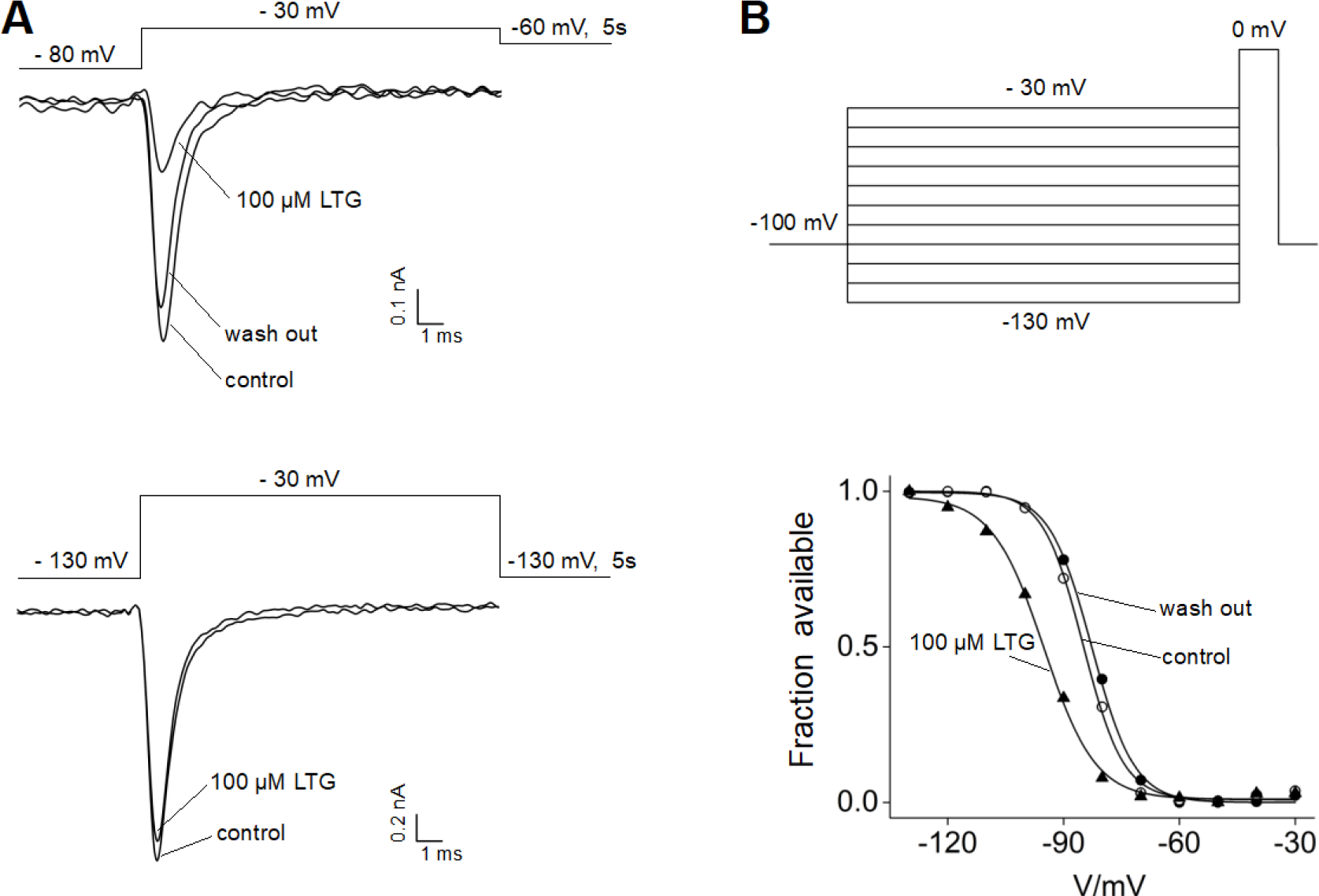
Effects of lamotrigine on Nav1.5 channels. A) The inhibition of Nav1.5 channel activity by lamotrigine (LTG) was tested at different holding potentials. Representative sodium currents recorded from HEK293T cells transiently expressing Nav1.5 channels before (control), upon 100μM LTG bath application (LTG) and after drug wash out (wash out) are shown. The cell was held at −80 mV (top) or –130 mV (bottom), then stepped to a test potential of –30 mV for 20 ms every 5 seconds. After the test potential the cell was stepped to –60 mV or –130 mV to keep the channel in the inactivated or resting state (top and bottom, respectively). Marked current inhibition can be noted in the traces shown on the top, consistent with the preferential binding of LTG to the inactivated state of sodium channels. B) LTG binding shifts the inactivation curve of Nav1.5 channel. Inactivating currents were elicited every 10 seconds using the voltage step protocol shown. The holding potential between each series of steps was set to –60 mV in order to keep the channel in the inactivated state. The current recorded at the test pulse of 0 mV was plotted against the voltage of the inactivating pulse to get the inactivation curve in control solution, upon bath application of 100 μM LTG, and after drug washout (traces from representative cell are shown at the bottom). Solid lines show fitting with a Boltzmann function, providing V1/2 values used to calculate the mean shift (ΔV, in mV): 13.6 ± 2.8 for 100 μM LTG and 2.1 ± 1.8 after drug washout (n=3). LTG treatment caused a leftward shift of the inactivation curve, rescued by the wash out of the molecule. Data represent mean ± S.E.M.

**Figure 2 – source data 1:**
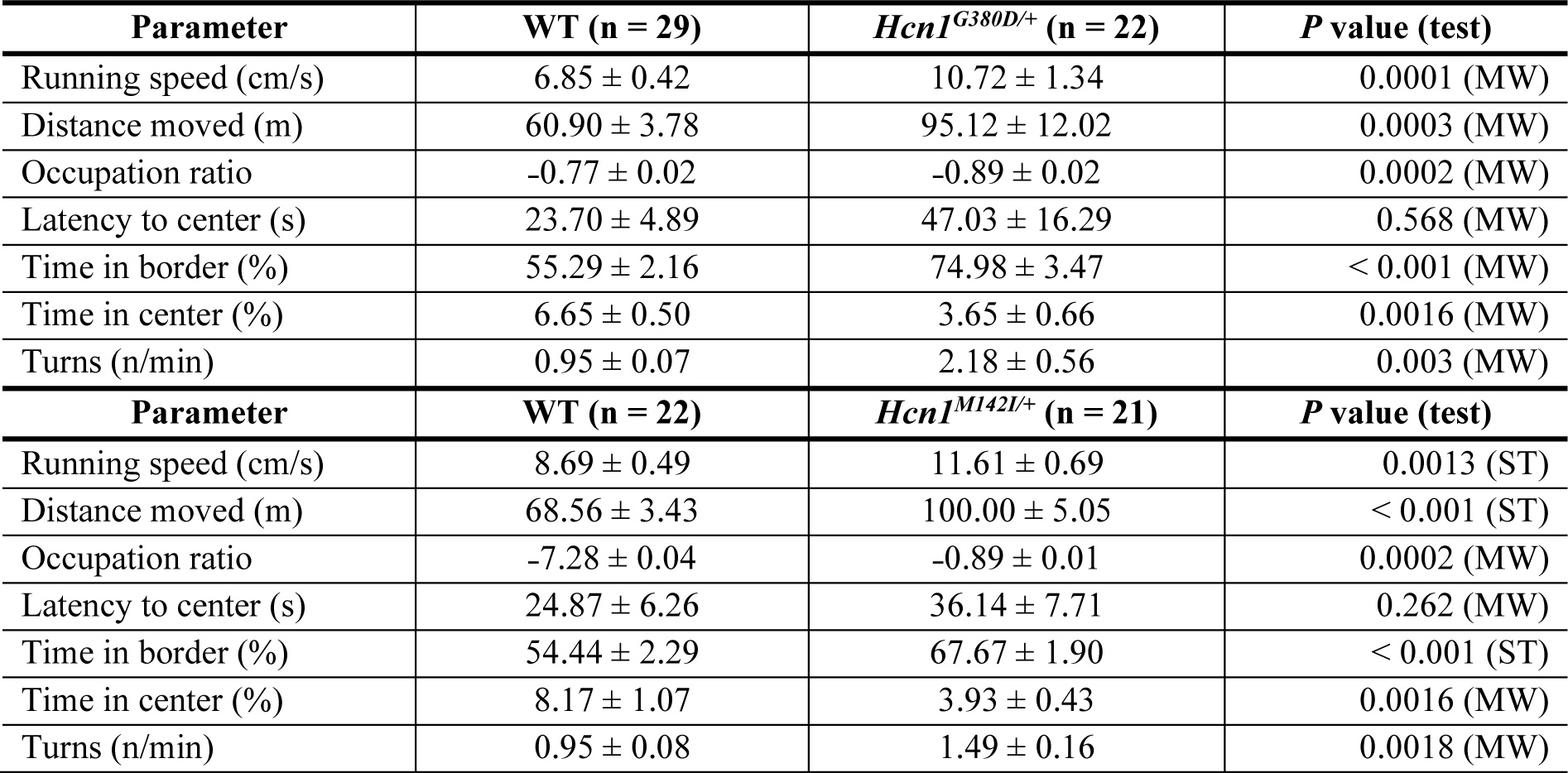
Open field parameter for Hcn1^G380D/+^ and Hcn1^M142I/+^ heterozygous mice in comparison to WT littermates. Times in border and center zones, respectively, are expressed as percentage of total trial duration. ST = Student’s t test, MW = Mann Whitney U test. Data represent mean ± S.E.M.

**Figure 2 – source data 2:**
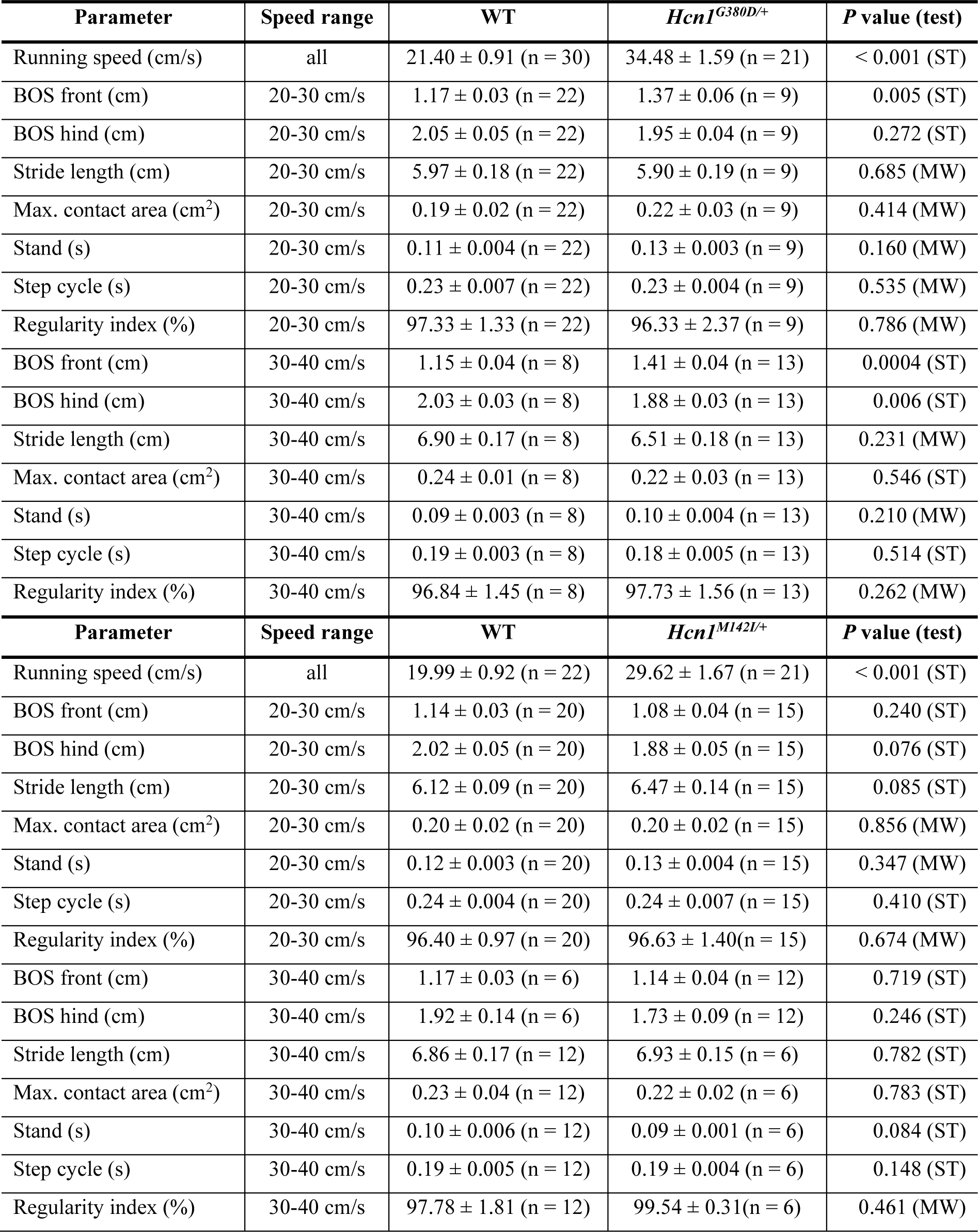
Catwalk gait analysis parameter for Hcn1^G380D/+^ and Hcn1^M142I/+^ heterozygous mice in comparison to WT littermates. Parameters listed correspond to the right front paw. Number of animals is shown in parenthesis. ST = Student’s t test, MW = Mann Whitney U test. Data represent mean ± S.E.M.

**Figure 8 – source data 1:**
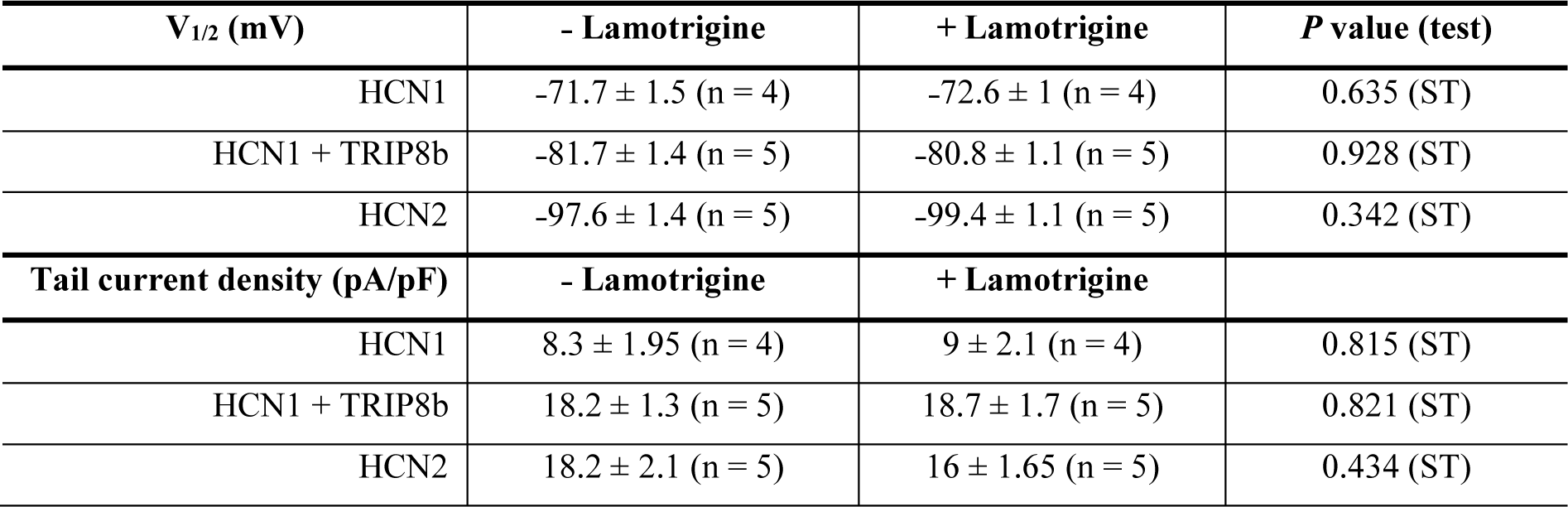
Lamotrigine has no direct effect on HCN1 or HCN2 channel activity. Number of cells is indicated in parenthesis. ST = Student’s t test. Data represent mean ± S.E.M.

**Figure 9 – source data 1:**
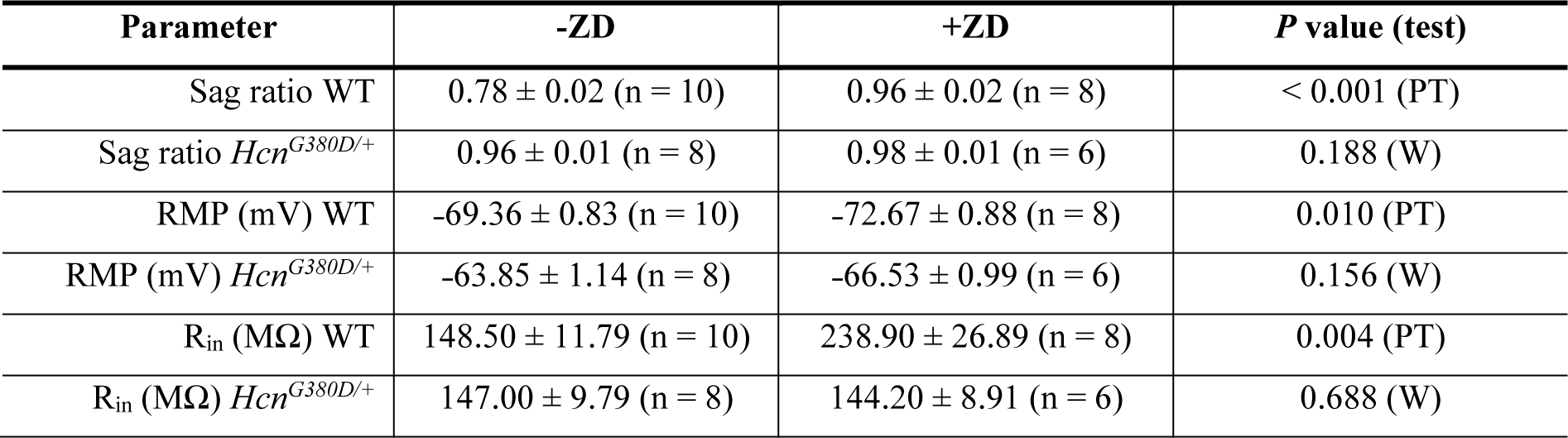
Lack of effect of ZD7288 on *Hcn1^G380D/+^* neurons. PT = paired *t* test, W = Wilcoxon matched-pairs signed rank test. Data represent mean ± S.E.M. Number of cells is shown in parenthesis.

